# A dispensable paralog of succinate dehydrogenase subunit C mediates standing resistance towards a subclass of SDHI fungicides in *Zymoseptoria tritici*

**DOI:** 10.1101/616904

**Authors:** Diana Steinhauer, Marie Salat, Regula Frey, Andreas Mosbach, Torsten Luksch, Dirk Balmer, Rasmus Hansen, Stephanie Widdison, Grace Logan, Robert A Dietrich, Gert HJ Kema, Stephane Bieri, Helge Sierotzki, Stefano FF Torriani, Gabriel Scalliet

**Author notes:** Kelly Scientific Resources, CH-4005 Basel, Switzerland. Novartis Pharma AG, CH-4002 Basel, Switzerland. Wellspring Biosciences, San Diego, CA 92121, USA. **Correspondence:** Gabriel Scalliet.

## Abstract

Succinate dehydrogenase inhibitor (SDHI) fungicides are widely used for the control of a broad range of fungal diseases. This has been the most rapidly expanding fungicide group in terms of new molecules discovered and introduced for agricultural use over the past fifteen years. A particular pattern of differential sensitivity (resistance) to a subclass of chemically-related SDHIs (SHA-SDHIs) was observed in naïve *Zymoseptoria tritici* populations. Class specific SHA-SDHI resistance was confirmed at the enzyme level but did not correlate with the genotypes of the succinate dehydrogenase (SDH) encoding genes. Mapping and characterization of the genetic factor responsible for standing SHA-SDHI resistance in natural field isolates identified a gene (*alt-SDHC*) encoding a paralog of the C subunit of succinate dehydrogenase. This paralog was not present within our sensitive reference isolates and found at variable frequencies within *Z. tritici* populations. Using reverse genetics, we showed that alt-SDHC associates with the three other SDH subunits leading to a fully functional enzyme and that a unique Qp-site residue within the alt-SDHC protein confers SHA-SDHI resistance. Enzymatic assays, computational modelling and docking simulations for the two types of SQR enzymes (alt-SDHC, SDHC) enabled us to describe protein-inhibitor interactions at an atomistic level and to propose rational explanations for differential potency and resistance across SHA-SDHIs. European *Z. tritici* populations displayed a presence (20-30%) / absence polymorphism of *alt-SDHC*, as well as differences in *alt-SDHC* expression levels and splicing efficiency. These polymorphisms have a strong impact on SHA-SDHI resistance phenotypes. Characterization of the *alt-SDHC* promoter in European *Z. tritici* populations suggest that transposon insertions are associated with the strongest resistance phenotypes. These results establish that a dispensable paralogous gene determines SHA-SDHIs fungicide resistance in natural populations of *Z. tritici*. This study paves the way to an increased awareness of the role of fungicidal target paralogs in resistance to fungicides and demonstrates the paramount importance of population genomics in fungicide discovery.

**Author Summary:** *Zymoseptoria tritici* is the causal agent of Septoria tritici leaf blotch (STB) of wheat, the most devastating disease for cereal production in Europe. Multiple succinate dehydrogenase inhibitor (SDHI) fungicides have been developed and introduced for the control of STB. We report the discovery and detailed characterization of a paralog of the C subunit of the SDH enzyme conferring standing resistance towards a particular chemical subclass of the SDHIs. The resistance gene is characterized by its presence/absence, expression and splicing polymorphisms which in turn affect resistance levels. The identified mechanism influenced the chemical optimization phase which led to the discovery of pydiflumetofen, exemplifying the importance of population genomics for discovery and rational design of the most adapted solutions.

## 1. Introduction

Fungicide research is driven by the discovery of molecules that either display novel modes of action or act on known targets but with a novel spectrum of biological activity or that escape target-based resistance mechanisms (1). During this research process, a very high diversity of molecules is generated to reach the necessary potency and biological spectrum. For single-site fungicides, rational active ingredient (AI) design and empirical chemical scouting are needed to best cover the chemical space of potential inhibitors. A broad biological spectrum is important for disease control. However, this is particularly difficult to achieve for single-site fungicides, mostly because the molecular targets usually show a significant level of variation across pathogens. In addition, the assessment of field populations’ sensitivity baselines and the definition of cross-resistance patterns are important since they may reveal unexpected variations which then need to be taken into consideration for AI design.

Such strategies were extensively used for the design of carboxamide Succinate Dehydrogenase Inhibitors (SDHIs). SDHIs block the tricarboxylic acid (TCA) cycle through inhibition of the succinate dehydrogenase enzyme (syn. succinate ubiquinone oxidoreductase (SQR), EC 1.3.5.1) which is better known as Complex II of the respiratory chain. SDHIs bind to the SQR enzyme at the ubiquinone binding site (Qp-site) which is created by the interface of three of the four enzyme subunits (2, 3). The fungal SQR is highly variable across species, mainly because of a low sequence conservation of the internal mitochondrial membrane SDHC and D subunits (4). These target variations have a big impact on the biological spectrum of activity of carboxamide SDHIs (5). Indeed, carboxin, the first molecule of this class introduced in 1966, displayed a basidiomycete spectrum of activity and was mostly used as seed treatment (6, 7). Major chemistry breakthroughs were needed to expand this biological spectrum to ascomycetes. In 2003, boscalid was released as the first foliar SDHI with a broadened spectrum of activity, enabling the control of diseases caused by ascomycetes (8). This discovery was shortly followed by the introduction of many other SDHIs covering almost the entire spectrum of fungal diseases (9). SDHI has been the fastest expanding class of fungicides in the past 15 years with 23 molecules currently listed by the fungicide resistance committee (10). In particular, some of these novel molecules effectively control the ascomycete *Zymoseptoria tritici* responsible for the main foliar disease of wheat. *Z. tritici* is the causal agent of Septoria tritici leaf blotch (STB), a major threat to bread and durum wheat production worldwide and a major driver for fungicide research (11). Resistance towards SDHIs was readily generated in the lab and caused by non-synonymous mutations within the Qp-site composing subunits encoded by *SDHB*, *SDHC*, and *SDHD* (12–14). Highly differential cross-resistance (XR) profiles were observed for some mutations. In particular, the SDHB_H267Y boscalid-resistant mutants showed increased sensitivity towards fluopyram in *Z. tritici* and multiple other species (13–16). The field situation is monitored by the industry and academic or governmental research institutes (17–19). To date, a panel of approximately 20 Qp-site subunit mutation types altering the activity of commercial SDHIs *in vivo* has been reported for *Z. tritici* populations in Europe (19, 20). The expected impact on field performance is variable depending on the particular mutation-SDHI compound combination (17). Overall for *Z. tritici*, the SDHI target resistance situation is at a stage of slight expansion in both diversity and frequency of mutations. The speed of resistance development and its practical impact on STB control has been contained, based on recommendations limiting the number of applications in spray programs and the use of mixtures with molecules carrying different modes of action.

Standing resistance towards fluopyram and isofetamid has been recently reported in *Z.tritici* European populations (21). The most shifted isolates were shown to display practical resistance to the compounds *in planta* but sensitivity to bixafen, another SDHI was not affected. Since no variation was observed in the sequences of the genes encoding the SQR Qp-site subunits, authors concluded that the mechanism was non-target based (21). During our research focusing on this novel class of SDHIs, we monitored the sensitivity baselines of a large collection of *Z.tritici* field isolates and identified similar resistance to fluopyram. This resistance was specific for a new chemical sub-class of SDHIs, which we termed SHA-SDHIs. Resistance was not associated with known mutations in SQR genes which was unexpected, since similarly to other fungicides used for STB control such as the QoIs (22–24) or the DMIs (25–27), previously known SDHIs resistance emerged through non-synonymous mutations within the target (17, 18). However, non-target related mechanisms have been reported that may contribute to sensitivity shifts, such as the overexpression of drug transporters like *MgMFS1* (28–31), or other transporters such as *ABCt-2* (32) or, a phenotypical connection between melanisation and fungicide uptake (33).

Therefore, the primary aim of this study was to characterize which molecular factors were involved in this natural SHA-SDHIs / fluopyram resistance. We report here the mapping and genetic validation of this resistance factor, a dispensable paralog of *SDHC* (*alt-SDHC*) which is present in 20-30% of the European *Z. tritici* population. Differential levels of expression and splicing of the alt-SDHC mRNA and a competition between the two SDHC proteins for inclusion into the SQR complex are the main factors modulating resistance. Molecular characterization of promoter sequences for a set of individuals revealed insertions of transposable elements in highly resistant isolates. This level of understanding enabled the careful design and early *in planta* assessment of pydiflumetofen, a novel SHA-SDHI affected by the mechanism but for which the variation has no practical impact on efficacy under normal use conditions. To our knowledge this is the first time that a fungicide target paralog with such complex presence/absence, splicing efficiency and expression polymorphisms has been described in naïve populations and taken into consideration during fungicide optimization.

## 2. Results

### 2.1. *Z. tritici* populations display differential sensitivity to the SHA-SDHI fluopyram

Assaying fungicide sensitivity of fungal populations is a pre-requisite for launching new fungicides. Large differences in sensitivity are frequently observed in naïve fungal populations which have not yet been in contact with the new fungicide. Depending on the fungicide and pathogen, the difference between least sensitive and most sensitive field isolates can reach a few orders of magnitude. Such differences are usually caused by standing variations either in the gene encoding the fungicide target or in its expression level. Variations in intracellular substrate abundance or expression of fungicide detoxification enzymes can also play a role in this differential sensitivity. Finally, cross-resistance plots help to determine whether similar factors affect different classes of chemicals with the same mode of action and to detect isolates that have been selected for their resistance to commercial fungicides with a mode of action similar to the newly released molecule. The early characterization of a fungal population’s sensitivity baselines is also an important tool for AI design, since it enables the early detection of potential standing resistance which would otherwise only become dominant when the new fungicides are tested in the field.

To observe whether similar factors affected the different classes of SDHIs, sensitivity towards commercial fungicides was determined for a set of 99 SDHI-naïve *Z. tritici* field isolates sampled in Europe between 2006 and 2009. The EC_50_ of these isolates was determined in liquid growth assays and the data obtained compared for each possible pair of SDHI fungicides (cross-resistance (XR) plots, Figure 1). As expected for fungicides carrying the same mode of action, a good correlation was observed for all SDHIs tested (Figure 1, ABCDEF). However, Spearman correlation factors were lower for fluopyram-paired comparisons (Figure 1). In particular, the three isolates displaying the lowest sensitivity (resistance) towards fluopyram (06STD024, 07STGB009 and 09STF011) displayed either normal or high sensitivity towards the other SDHIs benzovindiflupyr, boscalid and fluxapyroxad (Figure 1, ABC and DEF). The effect was specifically observed for fluopyram and all other research carboxamides carrying an aliphatic CC linker at the carbonyl end of the amide bond (data not shown, S1 Figure). Based on this observation, these molecules were grouped as a single cross-resistance group and termed stretch heterocycle amide SDHIs (SHA-SDHIs). *In vitro* enzyme succinate-quinone reductase (SQR) sensitivity tests were performed with mitochondria extracted from the highly shifted isolates, which indicated a target-based mechanism specific for SHA-SDHIs molecules (data not shown). However, in SHA-SDHIs/fluopyram shifted isolates, the target-related mechanism could not be explained by the genotypes of the known *SDHB*, *SDHC* and *SDHD* genes suggesting that other genes were involved in this fluopyram-specific resistance (S1 Dataset).

**Figure 1.**
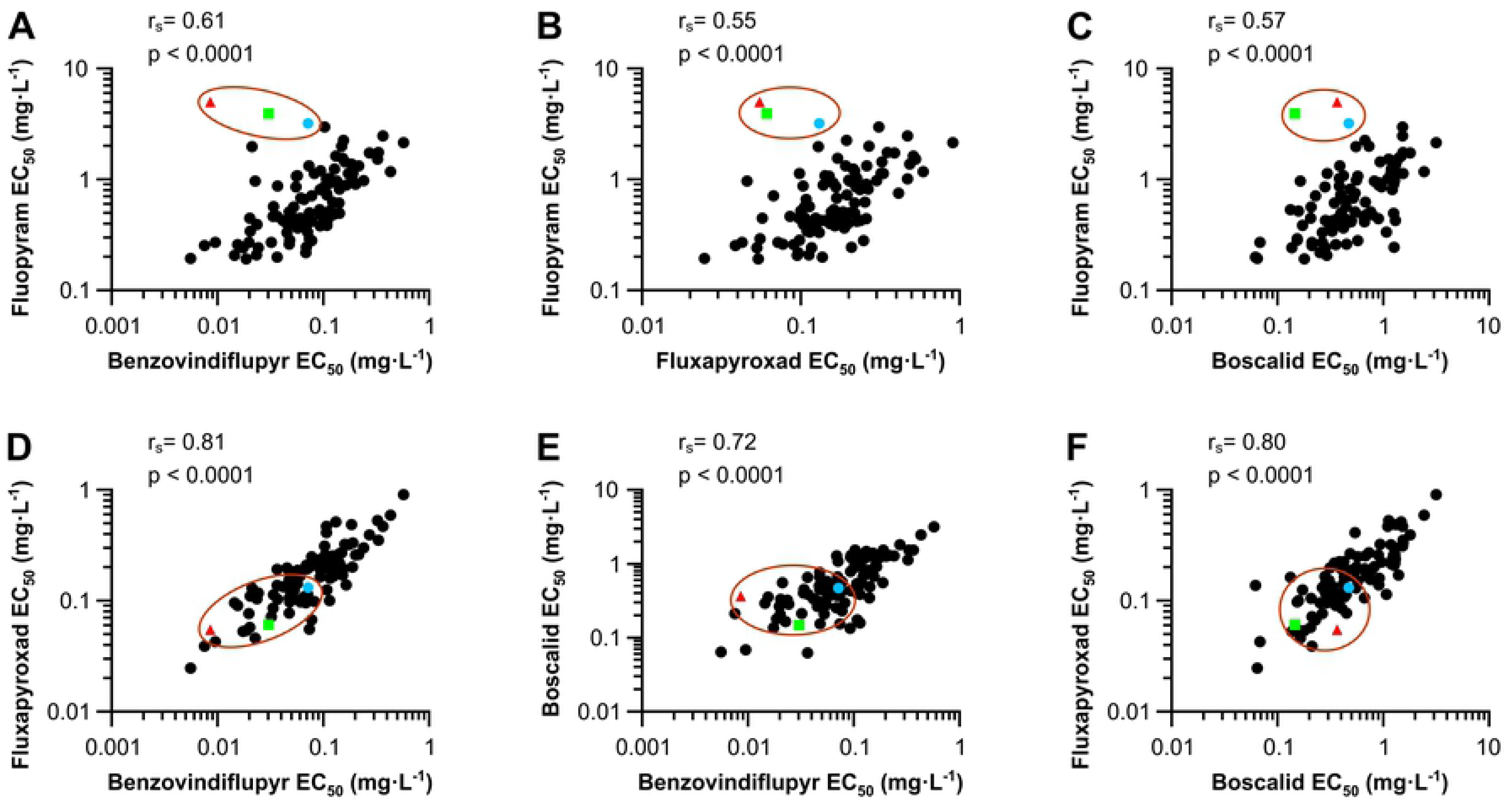
Baseline cross-resistance of Z. tritici populations to SDHI fungicides. Sensitivity towards different SDHIs was determined in liquid culture assays for a collection of 97 Z. tritici strains sampled for fungicide resistance monitoring in 2009 in Europe (plain circles). Two strains 06STD024 (red triangle) and 07STGB009 (green square), were considered fluopyram-resistant in monitoring performed in 2006 and 2007 respectively. 09STF011 (blue circle), belongs to the collection of 97 isolates sampled in 2009 and is the isolate with lowest sensitivity towards fluopyram in this set. Panels A, B and C represent liquid culture cross-resistance plots with SHA-SDHI fluopyram on the y axis and non-SHA SDHIs benzovindiflupyr, fluxapyroxad or boscalid on the x-axis respectively. D, E and F correspond to cross resistance plots of non-SHA SDHIs fluxapyroxad, benzovindiflupyr and boscalid, compared as pairs. 06STD024, 07STGB009 and 09STF011 are circled in red.

### 2.2. Mapping of a genetic factor responsible for fluopyram resistance in 06STD024 and 07STGB009 *Z. tritici* isolates

Crosses between *Z. tritici* isolates sensitive (S: IPO323, IPO94269) and resistant (R: 06STD024, 07STGB009) to fluopyram were generated. Mapping populations of 234 and 95 progeny were obtained for crosses IPO323 × 06STD024 and IPO94269 × 07STGB009 respectively. Progeny isolates from both crosses were characterized for their growth (R) / non growth (S) phenotypes on agar plates supplemented with 10 mg.L^−1^ fluopyram (Figure 2A). In both crosses, inheritance of the R phenotype was monogenic (49.5% and 51.5% resistant progeny respectively). A pooled sequencing bulked segregant analysis (BSA) approach was used to map the R locus using pools of genomic DNA from 30 S and 30 R progeny from cross IPO323 × 06STD024. Bulked segregant analysis (BSA) identified a locus on chromosome 3 between positions 3,081,782 and position 3,423,761 of IPO323 genome sequence (342kb) explaining the difference between the pools with 95% confidence (Figure 2B, S2 Dataset). Fine mapping with the full set of 234 IPO323 × 06STD024 progeny was performed with molecular markers such as cleaved amplified polymorphic sequences (CAPS) and direct PCR length polymorphisms developed from this region (S1 Table). This fine mapping located the resistance factor in an interval of 16kb from positions 3,200,730 to 3,217,341 of chromosome 3 of IPO323 (Figure 2B).

**Figure 2.**
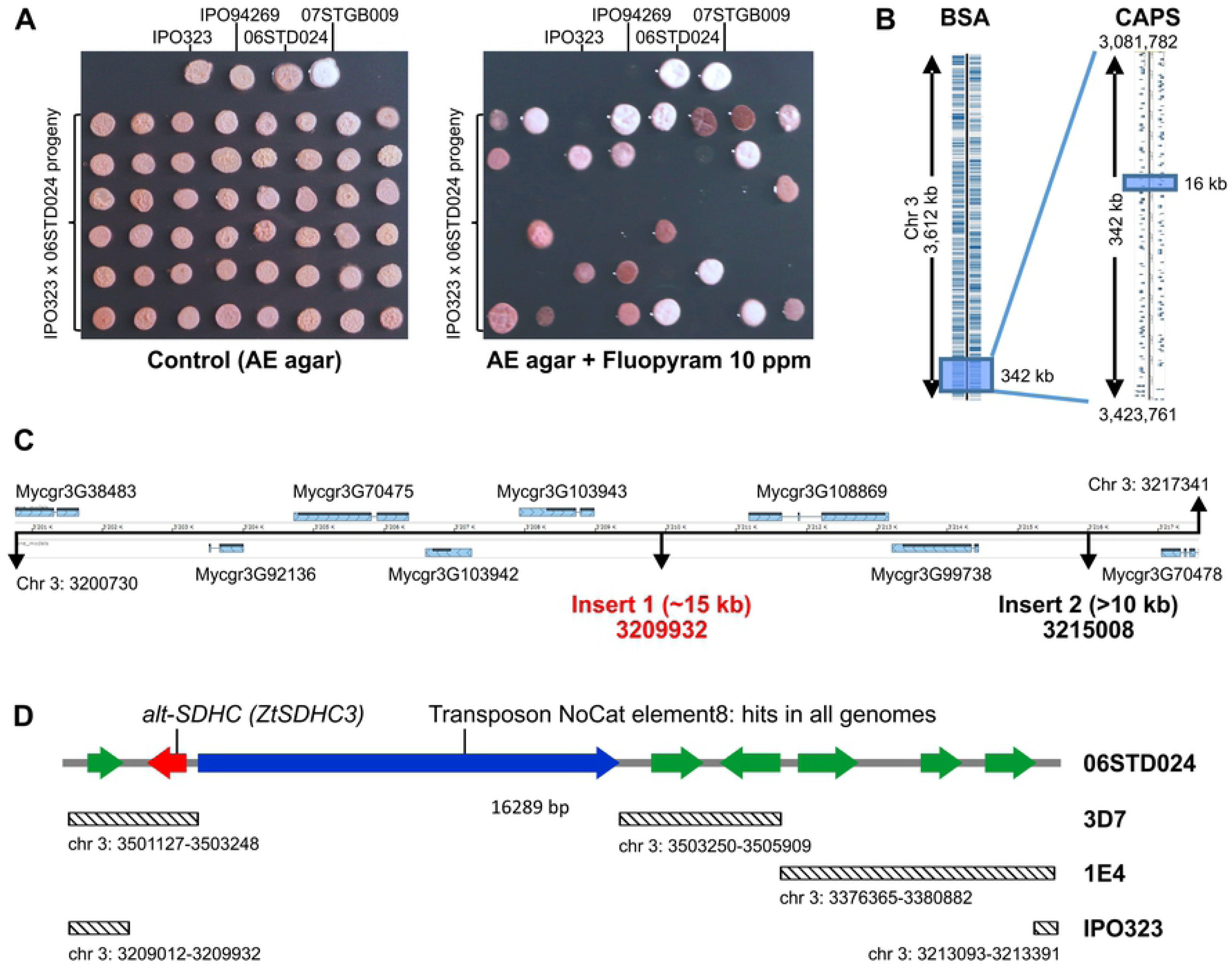
Fine mapping of fluopyram resistance factor using 06STD024 × IPO323 progeny. A. Agar plate growth assay used for characterizing progeny isolates for resistance or sensitivity to fluopyram. 2 µl of 2.10^6^ cells.ml^−1^ were spotted onto AE agar supplemented or not with 10 mg.L^−1^ fluopyram and incubated at 20°C. Pictures were taken either 7 days (control) or 14 days (fluopyram) after inoculation. B. IPO323 mapping intervals determined by BSA using 60 progeny isolates (i) and by CAPS markers (ii) on the full mapping population (234 progeny isolates). C. 16 kb mapping interval of IPO323 chromosome 3. Structural variations at this locus between IPO323 and 06STD024 were determined using long range PCRs. Insert 1 was fully sequenced, only borders of insert 2 were sequenced. Insert 1 and insert 2 positions are based on the IPO323 genome. D. Gene content of insert 1 region. Predicted genes and their orientation are visualized with arrows, green: putative genes, red: *alt-SDHC* (*ZtSDHC3*), blue: transposable element. Diagonally striped rectangles represent regions of high similarity (>90% identity) to other fully assembled *Z. tritici* genomes, corresponding chromosomal coordinates are indicated.

Within this genomic region, nine genes are predicted in IPO323 (Figure 2B). Only one gene, Mycgr3G70478 encoding a putative P-Type ATPase cation transporter, was predicted to be targeted to the mitochondria (S2 Table). Based on predicted function and subcellular localization, none of these genes could explain in simple terms the specific SHA-shifted SDHI sensitivity profile observed in the SQR enzymatic assay. Sliding-window PCRs were performed on 06STD024 genomic DNA to see whether structural variation may occur at the R locus that would potentially reveal additional genes in the resistant parental strain. This approach resulted in the detection of two large insertions at the mapped locus in the genome of the 06STD024 resistant strain that are not present in IPO323. The first large insertion was 15 kb in size and located at position 3: 3,209,932 of IPO323 genome. The second, over 10 kb in size was located at position 3: 3,215,008 of IPO323 genome (Figure 2C).

The 15 kb insert of 06STD024 was fully sequenced (GenBank: MK067274), and 7 putative CDS and a long putative transposon were identified within the locus (Figure 2D). One of these CDS displayed protein sequence similarity to SDHC (XM_003850403, 54% identity). The presence of two short introns within this CDS was confirmed by sequencing of 06STD024 cDNA and the corresponding gene was termed alternative SDHC (*alt-SDHC* or *ZtSDHC3*). Comparisons of the 06STD024-specific 15kb region of chromosome 3 to publicly available *Z. tritici* genomes identified similar regions in chromosome 3 of the 3D7 and 1E4 isolates (Fig 2D). The *alt*-*SDHC* gene was identified within the *Z. tritici* isolate 3D7 between positions 3: 3,502,409 and 3: 3,503,066 (GenBank: LT853694 locus tag ZT3D7_G4528 with 100% identity to *alt*-*SDHC* at the DNA sequence level). However, *alt-SDHC* was not identified in the 1E4 genome. Interestingly, the region of similarity between 06STD024 and 3D7 is interrupted by an uncategorized transposable element (TE) in 06STD024. This TE is 7 kb in length and located 182 bp upstream of the start codon of the *alt-SDHC* gene. The TE is found in multiple copies in IPO323 and within the other available *Z. tritici* genomes, but inserted at different chromosomal positions. *Alt-SDHC*-specific primers amplified this gene only in R parents 06STD024 and 07GB009, while *SDHC*-specific primers amplified the gene in all (R and S) parental strains. These PCR markers were used to genotype all progeny from crosses IPO323 × 06STD024 and IPO94269 × 07STGB009. For both crosses the presence of *alt*-*SDHC* fully segregated with the R phenotype (S1 and S3 Tables). Progeny from cross 07GB009 × IPO94269 were genotyped with additional CAPS markers from this chromosome 3 locus and confirmed the presence of the *alt-SDHC* gene at a similar chromosomal location in strain 07STGB009 compared to 06STD024 (S3 Table).

### 2.3. The alternative SDHC is responsible for fluopyram / SHA-SDHIs specific resistance

*alt*-*SDHC* (*ZtSDHC3*) DNA sequence displayed 62% identity compared to IPO323 *SDHC* (*ZtSDHC1*) CDS. The alt-SDHC protein sequence displayed an identity of 54% with IPO323 SDHC. The two nuclear encoded pre-proteins strongly differ at their N-termini. TargetP1.1 (34) predicted N-terminal mitochondrial transit peptides of 36 and 42 amino acids for alt-SDHC and SDHC respectively, that only share 16% identity. The predicted processed protein sequences (SQR cytochrome B subunit without transit peptide) of alt-SDHC and SDHC displayed a much higher similarity (62.5% identity). An alignment of the SDHC paralogs from *Z. tritici* is presented in Figure 3.

**Figure 3.**
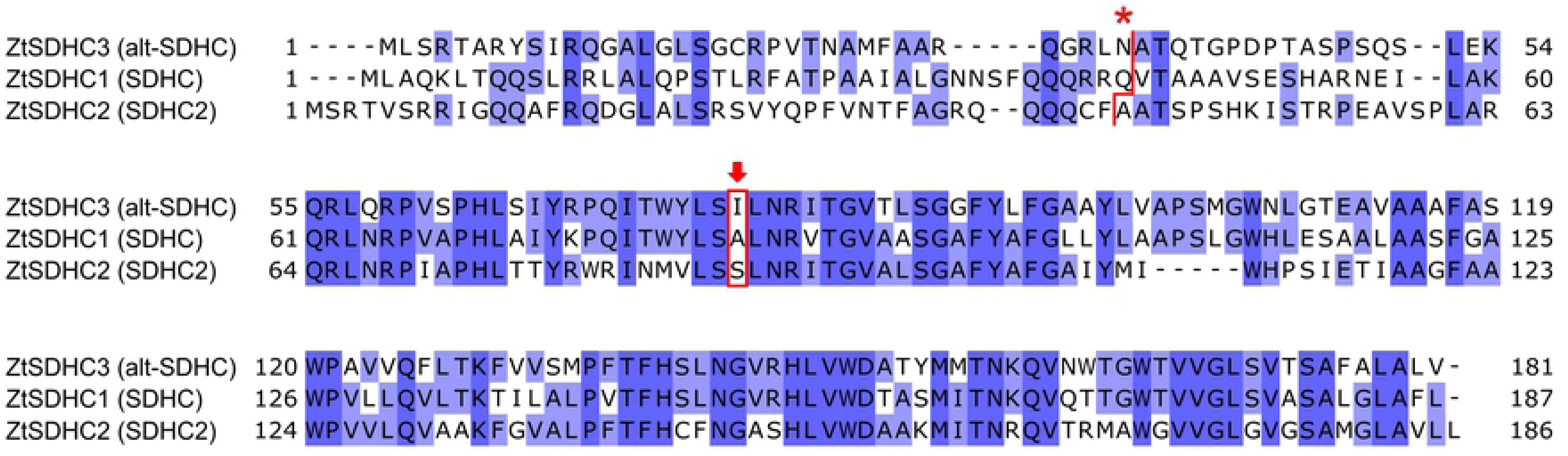
*Z*. *tritici* SDHC proteins alignment. *Z*. *tritici* ZtSDHC3 (alt-SDHC, NCBI MK067274, isolate 06STD024), ZtSDHC1 (SDHC, Uniprot F9XH52, isolate IPO323) and ZtSDHC2 (SDHC2, NCBI SMR59342, isolate IPO323) proteins were aligned with AlignX (Blosum62). Asterisk (*) is located above the predicted cleavage sites of the pre-proteins indicated by a red line. Red arrow highlights the Qp-site amino-acid residue likely involved in differential SDHI sensitivity pattern.

Phylogenetic analysis of fungal SDHC proteins revealed the presence of 0-2 paralogs in multiple species (S2 Figure). SDHC paralogs are found in multiple clades, but the number of paralogs within a genus appears species-specific (S4 Table). Another paralog of *ZtSDHC1* was identified in the IPO323 genome (Mycgr3G74581) and named *ZtSDHC2*. The Mycgr3G74581 gene model was modified using the revised gene model of Grandaubert *et al.* (35). This modified gene model was also found as a correctly predicted gene in the genome of isolate 1E4 (SMR59342). *ZtSDHC2* was present in all the genomes of sequenced *Z. tritici* isolates. Orthologs of *ZtSDHC2* were identified in *Z. brevis*, *Ramularia collo-cygni* and *Mycosphaerella emusae* genomes (S2 Figure). Orthologs of *ZtSDHC2* were not detected in the genomes of the closely related species *Pseudocercospora fijiensis*, *Dothistroma septosporum* and *Baudoinia panamericana*, which all carried an orthologue of *ZtSDHC1*.

This phylogenetic analysis suggested that *SDHC* was duplicated in a common ancestor of *Zymoseptoria spp.*, *Ramularia collo-cygni* and *Mycosphaerella emusae* to give *ZtSDHC2* and *ZtSDHC3*, the alternative SDHC. Species-specific losses of *ZtSDHC2* and/or *ZtSDHC3* must have occurred during the evolution of these species since *ZtSDHC3* is now only found in *Z. tritici*. The functional role of *ZtSDHC2* as a possible SQR C-subunit has not been validated. In the *Z. tritici* IPO323 isolate, there is no clear evidence of the expression of this gene in any tested condition (S3 Figure, (36)). Therefore, we concluded that *ZtSDHC1* is the only gene encoding a functional SDHC subunit in isolate IPO323, while isolate 06STD024 likely carries two functional SDHC subunits, SDHC encoded by *ZtSDHC1* and alt-SDHC encoded by *ZtSDHC3*.

To validate that *alt-SDHC* is responsible for fluopyram / SHA-SDHIs resistance, targeted deletions of *alt-SDHC* (*ZtSDHC3*) or *SDHC* (*ZtSDHC1*) were performed in the resistant isolate 06STD024. Targeted gene deletion vectors were constructed using a hygromycin resistance cassette flanked by 1-2kb of the upstream and downstream genomic sequences of either *alt-SDHC* or *SDHC* (see materials and methods). The *alt-SDHC* deletion mutants of 06STD024 were sensitive to fluopyram and other SHA-SDHIs. Their sensitivity levels were similar to IPO323, a SDHI-sensitive reference isolate (Figure 4A, 4B). The deletion of *SDHC* (*ZtSDHC1*) in 06STD024 was also achieved. These *SDHC* deletion mutants were more resistant (2 to 10 fold) to SHA-SDHIs than isolate 06STD024 (Figure 4B, S5 Table). The deletion of *SDHC* was not successful in IPO323 (data not shown), suggesting that *ZtSDHC2*, the unique *SDHC* paralog in this isolate, was not sufficient for maintaining SQR function in this background. IPO323 transformants carrying an ectopic insertion of a vector containing *alt-SDHC* under the control of a tetracyclin-repressible promoter were obtained (pTet::*altC*, Figure 4A). These IPO323 pTet::*altC* transformants displayed a SHA-SDHI resistance level similar or slightly superior to the 06STD024 isolate (Figure 4B, S5 Table). The addition of 30ppm doxycycline did not alter growth on non-selective media, but abolished growth in the presence of the SHA-SDHIs fluopyram and isofetamid (Figure 4A).

**Figure 4.**
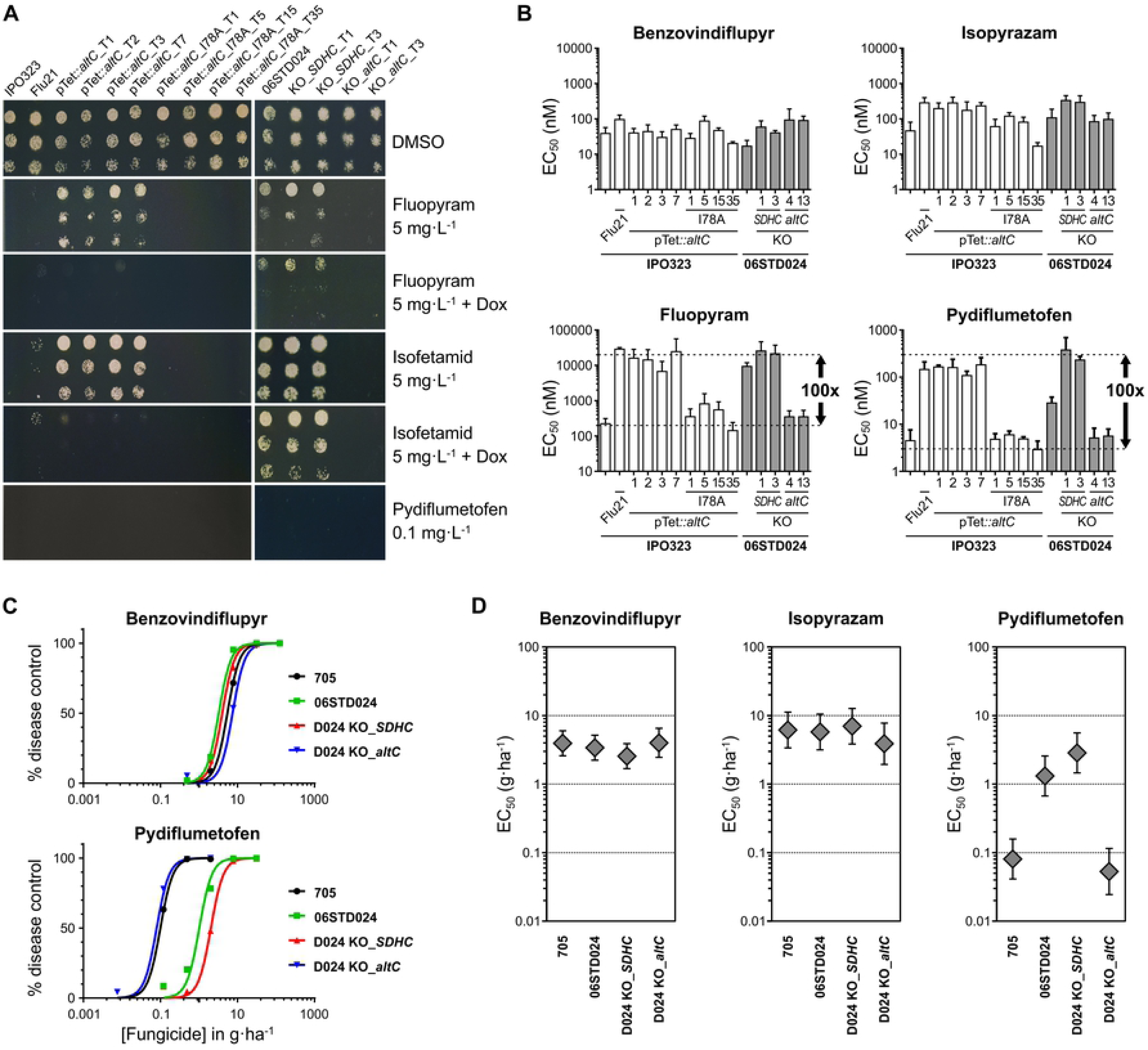
The role of alt-SDHC_I78 residue in conferring SHA-SDHIs-specific resistance *in vivo* and *in planta*. A. Agar growth phenotypes of IPO323 mutants (left panel) and 06STD024 mutants (right panel). Left panel: Flu21 is an IPO323 SDHC_A84I UV mutant, pTet::*altC* : IPO323 transformants carrying the *alt-SDHC* gene under the control of a tetracycline-repressible promoter, pTet::*altC*_I78A IPO323 transformants carry a similar construct but contain a mutated version of *alt*-*SDHC* gene encoding an I78A variant. Right panel: 06STD024 and individual deletion mutants of either the core *SDHC* (KO_*SDHC*) or the *alt*-*SDHC* (KO_*altC*). Pictures were taken at 6DPI, + Dox indicates the addition of doxycycline (30 mg.L^−1^) to the medium. B. Liquid culture sensitivity of IPO323 and 06STD024 mutants towards SDHIs. The set of characterized IPO323 (white bars) and 06STD024 (grey bars) mutants was similar to panel A. EC_50s_ (nM) were determined in duplicate in at least 3 biological replicates (see S5 Table). Values obtained for a broader range of marketed and research SDHIs are presented in S5 Table. C. *In planta* SDHI-sensitivity assays. The presented graphs are derived from a single biological experiment, each value / data point represents the mean disease control value of 12 individual plants. The sensitivity curves were obtained by non-linear regression of the data using GraphPad Prism software. D. *In planta* EC_50s_ (g.ha^−1^) of reference strain (705) and 06STD024 mutants for commercial SDHIs, benzovindiflupyr, isopyrazam (non SHA-SDHIs) and pydiflumetofen (SHA-SDHI). Values are derived from four biological replicates of 12 technical replicates each (EC_50_ +/− 95% confidence interval).

Amongst non-conserved positions in the protein alignment shown in Figure 3, isoleucine I78 of alt-SDHC corresponds to an alanine A84 in SDHC. A84 is located within the Qp-site, and is involved in ubiquinone substrate or inhibitor binding (14). Interestingly, the SDHC_A84I/V substitutions in *Z. tritici* were shown to confer resistance to fluopyram while displaying no effect on sensitivity/resistance to other SDHIs (14, 37). Therefore, the presence of an isoleucine at position 78 of alt-SDHC could explain the SHA-SDHIs-specific resistance profile conferred by the presence of *alt-SDHC*.

The involvement of the I78 Qp-site residue of alt-SDHC was tested by expressing an alt-SDHC_I78A variant in IPO323. IPO323 alt-SDHC_I78A transformants displayed similar sensitivity towards SHA-SDHIs as IPO323 or the 06STD024 *alt-SDHC* knock-out (KO) mutant (Figure 4A and 4B). Overall, these results demonstrated that *alt-SDHC* is responsible for the fluopyram/SHA-SDHIs-specific resistance profile of 06STD024 and that this gene can functionally replace *SDHC* in this background. *alt-SDHC* therefore encodes a dispensable functional C subunit of the *Z.tritici* SQR enzyme whose expression results in SHA-SDHIs specific resistance due to its natural I78 Qp-site residue.

The influence of the *alt-SDHC*-driven SHA-SDHI resistance for the control of *Z. tritici* during wheat infection was assessed with a small range of commercial SDHIs (Figure 4C, 4D). *In planta* SDHIs sensitivity assays were performed with the 06STD024 isolate and its *SDHC* or *alt-SDHC* KO mutants. A control strain (705) devoid of the *alt-SDHC* gene but more aggressive than IPO323 on wheat variety Riband was also included for comparison (Figure 4C and 4D).

On untreated plants, 06STD024 KOs displayed infection levels similar to wild type, although a slightly delayed virulence was observed for the *SDHC* KO (data not shown). *In planta* sensitivities towards the non SHA-SDHIs benzovindiflupyr and isopyrazam were similar across the isolates (Figure 4D). Conversely, the presence of *alt-SDHC* impacted the SHA-SDHI compound pydiflumetofen (Figure 4C, 4D). Similarly to liquid culture assays, 06STD024 and its *SDHC* or *alt-SDHC* KO mutants differed in their sensitivity towards the SHA-SDHI pydiflumetofen. The most shifted isolate was 06STD024 *SDHC* KO, which displayed an *in planta* EC_50_ 53 fold higher than 06STD024 *alt-SDHC* KO mutant (2.85 g.ha^−1^ and 0.053 g.ha^−1^ respectively) (Figure 4D). Isolate 06STD024 displayed a reduced *in planta* EC_50_ of 1.31 g.ha^−1^ which corresponds to a sensitivity difference of 25 fold compared to the *alt-SDHC* KO mutant.

Our data validate the effect of *alt-SDHC* on *Z.tritici* sensitivity towards commercial SHA-SDHIs *in planta*. The activity of pydiflumetofen on the most SHA-SDHI-shifted *Z. tritici* GM isolate was similar to that of benzovindiflupyr on wild type isolates. The strongly shifted *alt-SDHC* genotypes are therefore not considered to be resistant to the compound in practice.

### 2.4. Expression levels of the two types of SDHC subunits influence mitochondrial SQR composition and resistance

In order to explore the influence of differential alt-SDHC expression on SQR enzyme composition and resistance, we used reference isolates 06STD024, IPO323 and the pTet::*altC* IPO323 transformant grown under inductive or repressive conditions and characterized i) *SDHC* and *alt-SDHC* mRNA expression ii) mitochondrial SDHC and alt-SDHC proteins abundances (quantified by LC-MS/MS) and iii) SQR enzyme sensitivity to SDHIs (Figure 5, Table 1).

**Figure 5.**
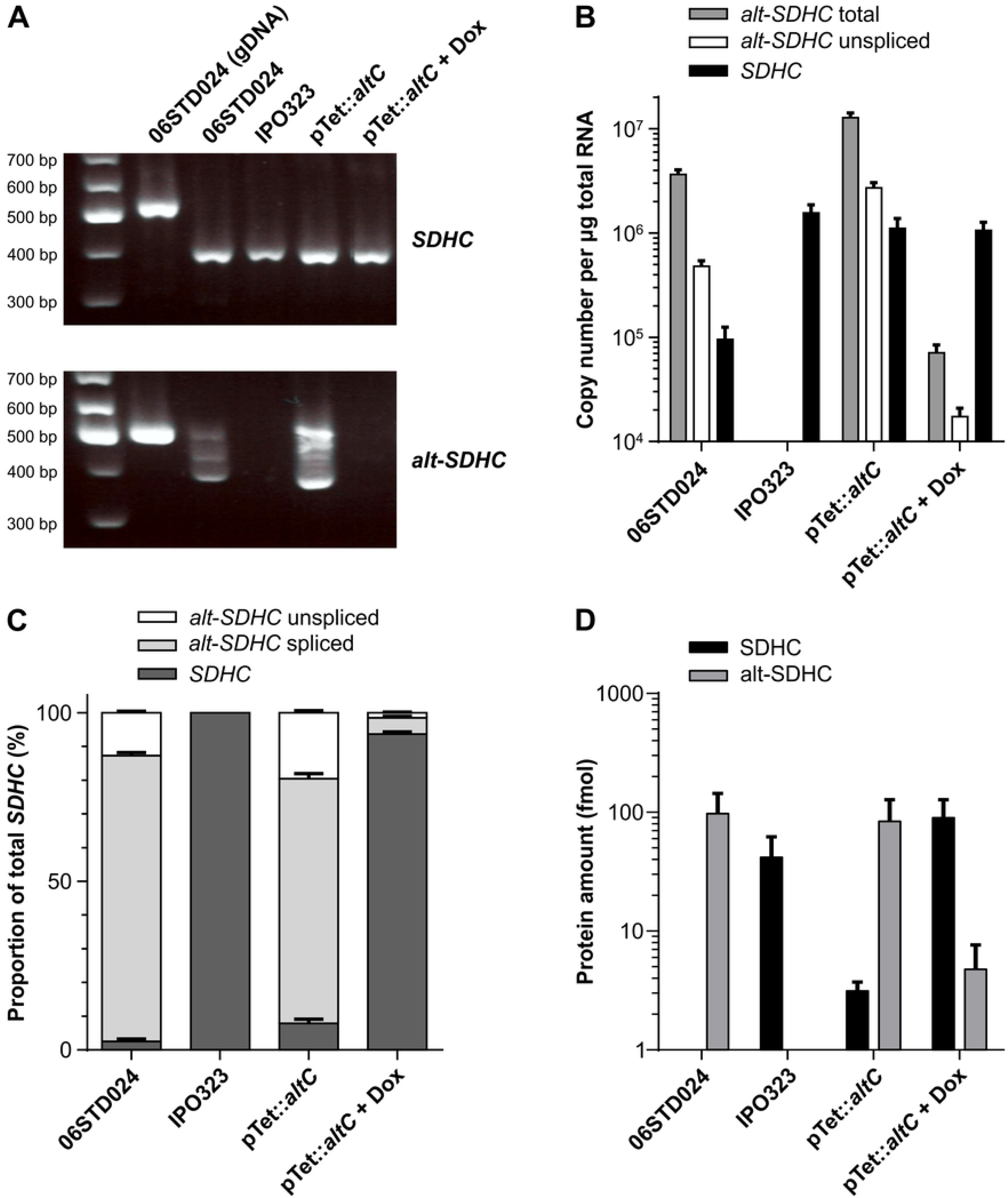
Expression-driven competition of SDHC and alt-SDHC proteins for functional integration in the mitochondrial SQR. A. RT-PCR analysis of *SDHC* and *alt-SDHC* in 06STD024 and IPO323 pTet:*altC* transformant. The expected PCR products corresponding to fully spliced mRNAs were 389 and 384 bp for *SDHC* and *altSDHC* respectively. B. Absolute quantification by RT-qPCR of the three *SDHC* mRNA species in the 06STD024 strain and IPO323 pTet::*altC* transformant. C. Normalized proportion of the three mRNA species (as deducted from panel B). D. LC-MS/MS quantification of the SDHC and altSDHC proteins in mitochondrial extracts from 06STD024 and IPO323 pTet::*altC* transformant. Values presented are the mean of 6 individual experiments ± SD.

**Table 1.**
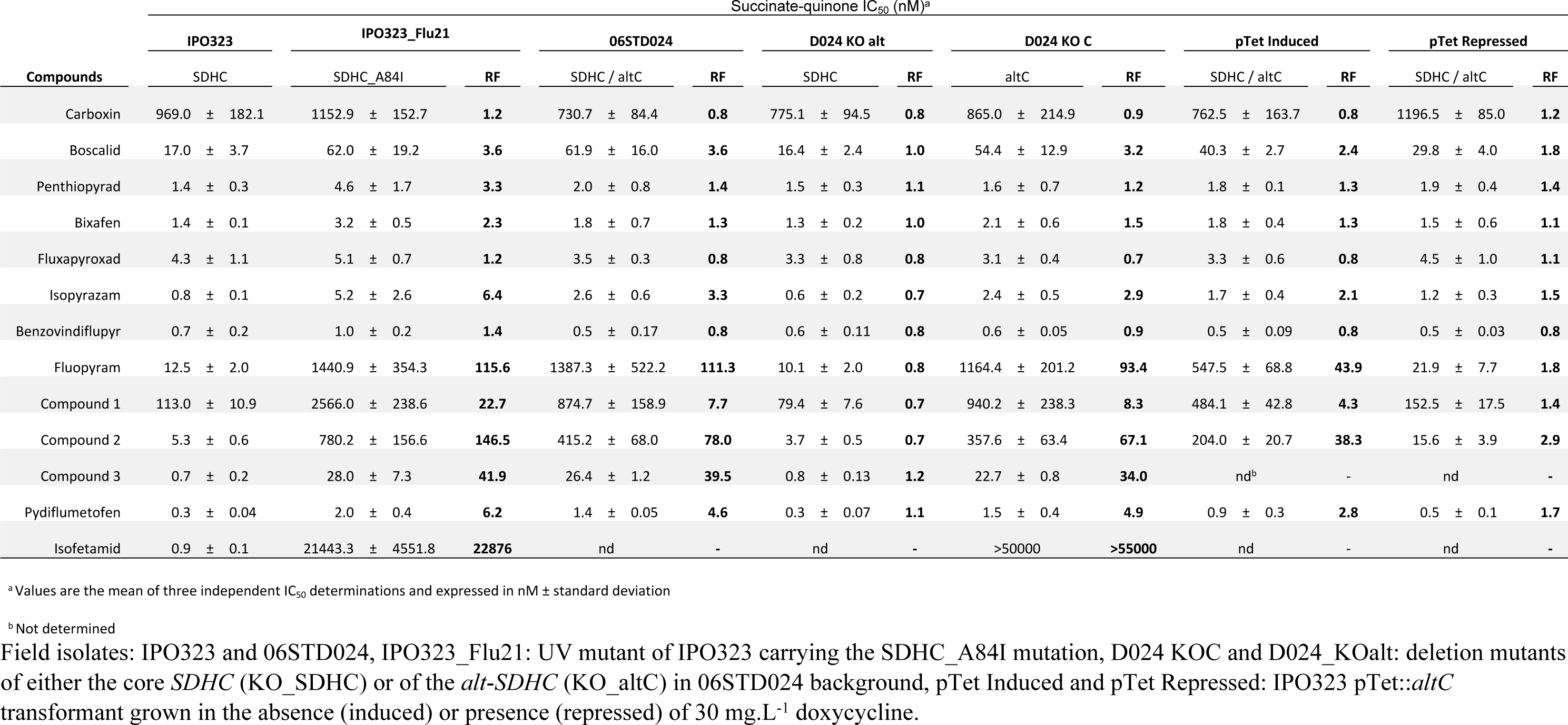
Succinate-quinone SDHIs sensitivity assays on purified mitochondria of field isolates and transformants of *Z. tritici*.

RT-PCR revealed partially incomplete splicing of *alt-SDHC* mRNA in 06STD024. A band of non-spliced mRNA of a size similar to the genomic amplicon was clearly visible as well as other partially spliced species of higher size compared to the main fully spliced band (Figure 5A). This partial splicing was not only observed with 06STD024 but also with the ectopic transformant of IPO323 expressing the *alt-SDHC* gene under the control of the tetracyclin-repressible promoter (Figure 5A). Conversely, the *SDHC* gene appeared to be fully spliced as suggested by a single band of the expected size (Figure 5A). Hydrolysis probe RT-qPCR assays were used to quantify unspliced (second intron) and total forms (third exon) of *alt-SDHC* mRNA as well as the total form (spliced) of *SDHC* mRNA (third exon). These RT-qPCR assays enabled the comparison of functional mRNA quantities and ratios for both *SDHC* genes (Figure 5B, 5C).

The total amount of *SDHC* mRNA was ten folds lower in 06STD024 compared to IPO323, demonstrating strain to strain variation (Figure 5B). As expected in the pTet::*altC* IPO323 transformant *alt-SDHC* mRNA was strongly induced in the absence of doxycycline and highly repressed by doxycycline 30ppm (100 fold). This differential *alt-SDHC* mRNA expression had no impact on *SDHC* expression (Figure 5B). In 06STD024, spliced *alt-SDHC* mRNA was 34 fold more abundant than *SDHC* mRNA (84.6% spliced *alt-SDHC* compared with 2.5% spliced *SDHC*). In this context, the alt-SDHC protein was the only SDHC protein detected in the 06STD024 mitochondrial sample (97 fmol). In the IPO323 pTet::*altC* transformant grown under non-repressive conditions, fully spliced *alt-SDHC* mRNA was nine fold more abundant than the *SDHC* mRNA (72.5% spliced *alt-SDHC* compared with 7.9% *SDHC*) (Figure 5C). In this context, the mitochondrial alt-SDHC protein was 27 fold more abundant than the SDHC protein (84 fmol alt-SDHC vs 3.1 fmol SDHC, Figure 5D). Adding 30 ppm doxycycline repressed the expression of *alt-SDHC* (Figure 5A, B) which resulted in 20 fold lower abundance compared to *SDHC* mRNA (93.7% of SDHC mRNA compared to 4.7% spliced *alt-SDHC* mRNA, Figure 5C). In this context the SDHC protein dominated over the alt-SDHC protein in the mitochondria (90 fmol SDHC vs 4.8 fmol alt-SDHC, Figure 5D).

Although the amounts of spliced mRNA encoding the alt-SDHC protein differ significantly in the pTet::*altC* IPO323 transformant grown under permissive versus repressive conditions, the total amount of mitochondrial SDHC subunit was similar (87.1 fmol vs 94.8 fmol) suggesting a saturation limit caused by the availability of other SQR subunits for integration into the functional SQR complex *in vivo*. In this context, the competition between SDHC and alt-SDHC proteins for integration into the SQR enzyme also translates into a steep reduction of the least expressed subunit as found for SDHC in 06STD024.

The impact of these mixed SDHC compositions on SQR enzyme sensitivity towards SHA-SDHIs was tested using succinate-ubiquinone enzyme inhibition tests (Table 1). Mitochondrial SQR from a non-repressed IPO323 pTet::*altC* transformant displayed IC_50_ values clearly shifted for SHA-SDHIs (RF fluopyram = 44) but this resistance level was lower than 06STD024 (RF fluopyram = 111) (Table 1). Conversely IC_50_ values obtained with mitochondria extracted from the same transformant grown in the presence of doxycycline displayed very low resistance to SHA-SDHIs (RF fluopyram = 1.7) and IC_50_ values similar to sensitive isolates IPO323 or 06STD024 KO_altC (Table 1).

These results are consistent with a mixture of the two types of SQR enzymes being simultaneously present and functional. They suggest a competition for integration within the functional enzyme leading to the presence of mixed SQR populations and mitigating the observed sensitivity shift due to differing expression ratios of the two types of C subunits.

### 2.5. Molecular docking within 3D models of SQR variants explain differential potency and cross resistance among SHA-SDHIs

Significant potency and sensitivity variations have been observed for the molecules that were tested against *Z*. *tritici* SQR variants (Table 1). Amongst the SHA-SDHIs molecules tested, pydiflumetofen combined highest potency on wild type SQR (IC_50_=0.3nM) with lowest resistance levels on alt-SQR (<7). In contrast, isofetamid showed a dramatic loss of efficacy on both alt–SQR and C_A84I-SQR mutants (RF>20’000) despite a high potency on WT-SQR (IC_50_=0.9mM). Finally, fluopyram combined moderate potency (IC_50_=12.5nM) on WT-SQR with an approximate 100 fold resistance on the alt-SQR. To unravel the factors driving potency and resistance across SHA-SDHIs at an atomistic level, 3D homology models for the WT-SQR and alt-SQR were generated and comparative docking studies carried out.

The superposition of WT and alternative SQR models showed that the two enzymes are structurally highly conserved. In particular all Qp site residues are conserved except SDHC_A84 which corresponds to I78 in alt-SDHC (Figure 3, Figure 6A). In agreement with this, C_A84I and alt-SQR enzymes displayed highly similar SDHI sensitivity profiles (Table 1). Molecular docking of SHA-SDHIs into the homology models of the *Z.tritici* SQR variants have been carried out and protein-ligand interactions analyzed. The interaction of carboxamide SDHIs with the SDHC_A84 residue and C_A84V/I-SQR mutants has been described previously (14). Carboxamide SDHIs are predicted to interact with SDHC_A84 via Van-der-Waals forces and a change from alanine to a larger valine or isoleucine residue is therefore likely to have an impact on ligand binding. This impact was specifically observed for fluopyram which was affected by high resistance factors compared to carboxin, boscalid, and isopyrazam in SDHC_A84V/I UV mutants. We assumed that this was caused by the linker of fluopyram which is in the z-dimension sterically more demanding and cannot be properly accommodated with a valine or an isoleucine at position 84 (14). These findings and assumptions remain true for fluopyram interaction with the alt-SQR.

**Figure 6.**
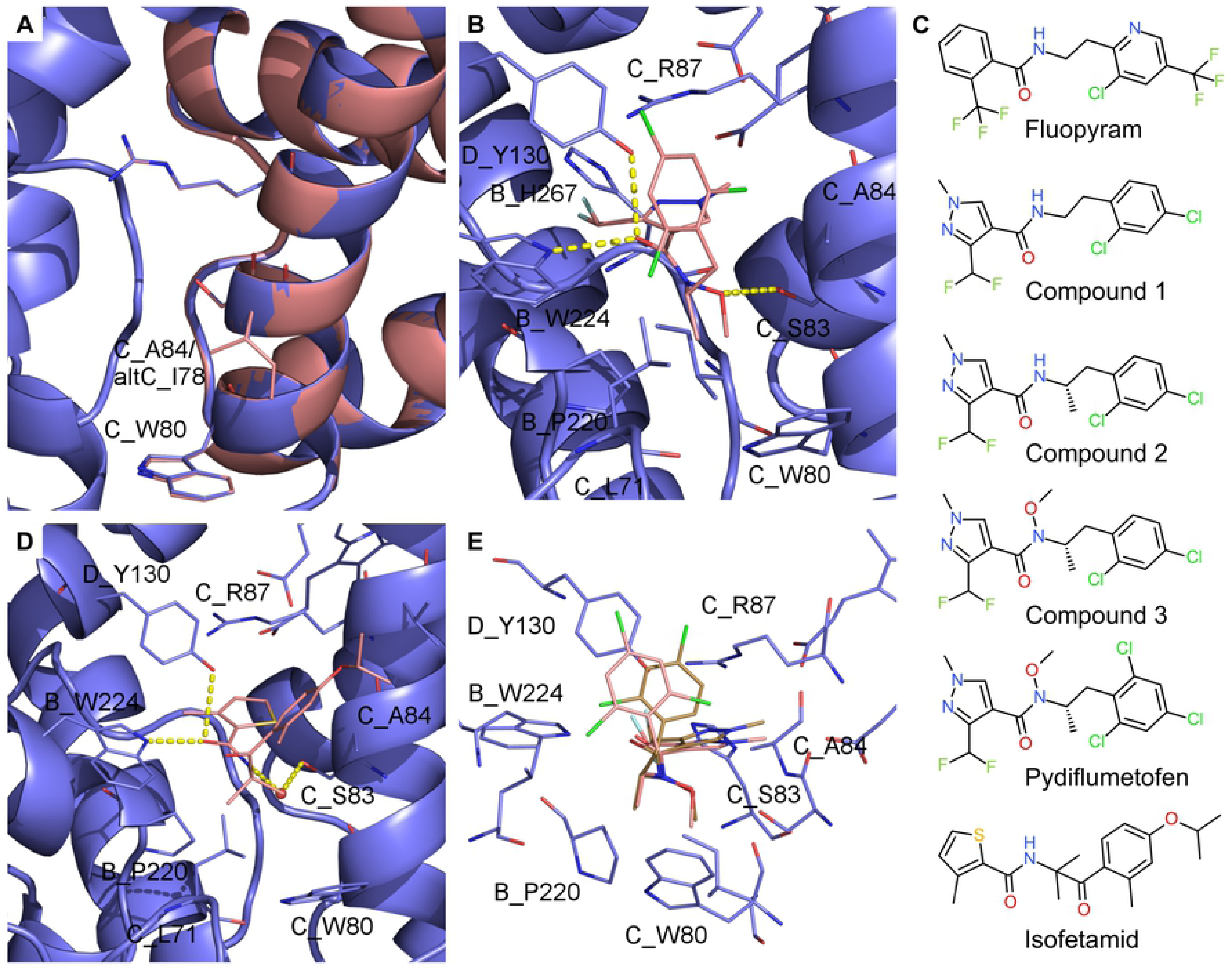
Comparison of *Z. tritici* 3D models of the two SQR paralogs and putative binding modes for SHA SDHIs. A. Homology model of *Z. tritici* WT-SQR (blue) superimposed on the homology model of *Z. tritici* alt-SQR (salmon). B. Putative binding mode of pydiflumetofen in *Z. tritici* WT-SQR. C. 2D depiction of SDH inhibitors used for docking comparisons and discussed in the text. D. Putative binding mode of isofetamid in *Z. tritici* WT-SQR. E. superposition of energy minimum conformations for pydiflumetofen and compound 3.

Molecular docking of isofetamid into the WT-SQR predicts hydrogen bonds between the amide oxygen of the molecule and SDHD_Y130 and SDHB_W224 residues (Figure 6D). The nitrogen of the amide forms a hydrogen bond to SDHC_S83 mediated by a water molecule. The carbonyl oxygen of isofetamid SHA aliphatic chain is not involved in a hydrogen bond to SQR but plays an important role together with the gem di-methyl group for the pre-organization of the molecule into the bioactive conformation. In particular, the ortho methyl substituent stabilizes a conformation of the phenyl ring in which Van-der-Waals interactions to SDHC_A84 can be formed. In addition, the para isopropyloxy substituent is at the right distance in the model to form Van-der-Waals interactions to SDHC_V88. Contrastingly, in alt-SQR the isoleucine 78 of alt-SDHC reduces the size of the binding pocket. Docking of isofetamid into the smaller binding pocket of alt-SQR did not result in any energetically favorable conformation which is in agreement with the very poor potency of isofetamid on alt-SQR in enzymatic tests (Table 1). Maintaining an isofetamid conformation similar to the one obtained in WT-SQR leads to a steric clash of the phenyl ring with alt-SDHC_I78 which is in agreement with the very high resistance factors (>20’000 fold) observed in the alt-SQR and A84I-SQR mutants.

In contrast, pydiflumetofen is highly potent on WT-SQR (EC_50_=0.3nM) and is only shifted by a factor of 6 in the isoleucine SQRs (C_A84I and alt-SQR) (IC_50_=2.0 nM). This is a unique behavior to our knowledge within SHA-SDHIs. To assign how distinct parts of the chemical structure of pydiflumetofen contribute to the favorable activity and resistance profile, molecules have been selected for analysis that belong to the same chemical series of pydiflumetofen but differ only by single chemical transformations (matched pairs, compounds 1-3 shown Figure 6C). The putative binding mode of pydiflumetofen in complex with the classical and alternative SQR are shown in Figure 6 B, E.

Similarly to other carboxamide SDHIs, the binding interaction of pydiflumetofen with the Qp site involves hydrogen bonds with SDHD_Y130 and SDHB_W224 through the carbonyl oxygen of the amide bound. The specific SHA-SDHI CC linker of pydiflumetofen bears a stereo center. The role of this stereo center is elucidated by comparing two related SDH inhibitors for which the only difference is a methyl group: compound 1, a molecule bearing an ethyl linker without methyl group and stereo center, is more flexible and multiple low energy conformations exist. A pre-organization is caused by the additional methyl group in compound 2, which also introduces a stereo center (R and S enantiomers). The S enantiomer is predicted to adopt a low energy conformation more compatible with the shape of the ubiquinone binding pocket. This pre-organized conformation leads to a favorable entropic effect that is predicted to increase the activity. The IC_50_ for compound 2 is indeed lower in comparison to compound 1 on WT-SQR but the magnitude of the effect (21 fold) is more pronounced than predicted. The same trend is observed in isoleucine SQR but in this case leading to only a 3 fold increased activity for compound 2 compared to compound 1.

A very special feature of pydiflumetofen is its substituted N-methoxy amide. While all other carboxamide SDHIs are predicted to form hydrogen bonds to SDHC_S83 mediated by a water molecule, pydiflumetofen is predicted to form a direct hydrogen bond of the methoxy oxygen to the serine (Figure 6B). This hypothesis is in line with various crystal structures in which ubiquinone analogues are bound to SQR (e.g. pdb code 5C3J), and form direct hydrogen bonds to the serine. In addition, in the model the methyl moiety of the N-methoxy amide forms lipophilic interactions with isoleucine 269 and proline 220 of SDHB. This might be the reason for the 7.6 fold increased potency of N-methoxy amide-containing compound 3 compared to the matched pair compound 2 (without N-methoxy amide). The resistance factors are reduced to 2 or 39.5 fold for compound 3 in comparison to 146 or 80 fold for compound 2 which is hypothesized to be due to positive lipophilic interactions to I84 or I78 in the SDHC or alt-SDHC SQR variants respectively.

Interestingly the addition of a third chlorine atom in the aromatic ortho position significantly decreases the resistance factor from 42 fold for compound 3 to 6 fold for pydiflumetofen. A conformational analysis showed that the aromatic ring is rotated further away from SDHC_A84 in comparison to compound 3 in the energy minimum conformation of pydiflumetofen (Figure 6E). It is assumed that this particular conformational effect reduces the steric hindrance in alt-SQR.

### 2.6. Polymorphism of *alt-SDHC* in *Z. tritici* field populations: presence/absence, expression and splicing

The presence/absence polymorphism of *alt-SDHC* and *SDHC* in *Z. tritici* field populations was determined using PCR specific for each gene (see materials and methods). The *alt-SDHC* gene was detected at frequencies ranging from 17% to 31% in the EU depending on the year of sampling whereas the *SDHC* gene was detected in all isolates (Table 2). 123 *Z. tritici* genomes (38) corresponding to isolates collected in four locations (Switzerland, USA, Israel, Australia) prior to the introduction of SDHIs for disease control in wheat were screened *in silico.* The *alt-SDHC* gene was found in 29% of Swiss isolates and in 18% of the USA (Oregon) isolates. Interestingly, the gene was not detected in the 25 isolates from Israel but was present in all isolates from Australia.

**Table 2.**
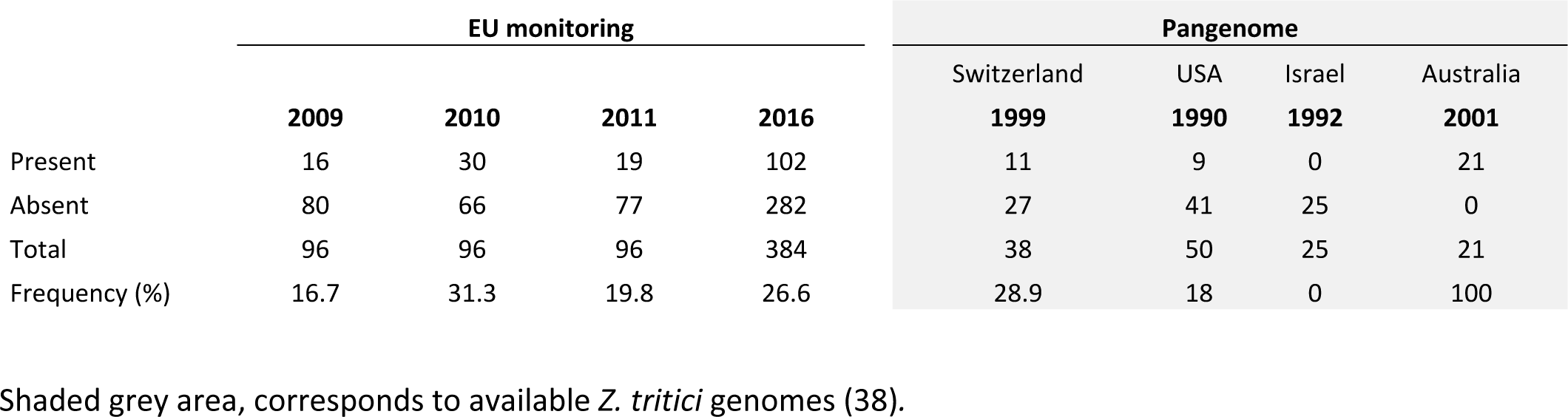
Frequency of alt-SDHC in *Z. tritici* populations. Occurrence of the *alt-SDHC* gene in European monitoring populations and the pangenome.

*alt-SDHC* sequences were determined by Sanger or Illumina amplicon sequencing for a panel of 154 isolates carrying the gene (EU collections from Table 2). We identified 12 nucleotide haplotypes (S6 Table) among which 11, were rare variants of the main canonical sequence and represented only once in the panel (0.6%). Among these variants, six carried non-synonymous mutations affecting the alt-SDHC protein sequence. Three corresponded to truncated likely inactive forms of the protein and three corresponded to likely functional R34Q, S66Y and T73S protein variants. In comparison, for 350 strains (2016 collection) sequenced at the *ZtSDHC1* gene locus, 206 different nucleotide haplotypes were identified for a total of 27 different protein variants (data not shown). This relatively rare occurrence of mutations within the *alt-SDHC* gene is highly contrasting with the high degree of polymorphisms observed for the core *SDHC* gene.

Liquid culture and plate growth SDHI sensitivity assays were performed on a cohort of 93 field isolates collected in 2009 and characterized for the presence/absence of the *alt-SDHC* gene (Figure 7A, 7B). Liquid culture assay validated a significant difference between the two groups with SHA-SDHI fluopyram (t test, p<0.05) but not with non SHA-SDHI benzovindiflupyr (Figure 7A). However, the panel of alt-SDHC containing isolates displayed a wide range of fluopyram EC_50s_ varying from sensitive (0.3 mg.L^−1^) to resistant (up to 3.2 mg.L^−1^). The growth/no growth phenotype on SHA-SDHI supplemented agar plates of these 93 field isolates mostly correlated with the presence/absence of the *alternative SDHC* gene (Figure 7B). However, again depending on the SHA-SDHI used for the assay, significant growth/sensitivity differences are visible across the isolates carrying the alt-SDHC gene. Among SHA-SDHIs, isofetamid was the compound for which the presence of the gene gave the clearest correlation (presence=growth, absence=no growth). Only one isolate carrying the gene, 09STIR20.1 did not grow on isofetamid-supplemented agar plate (10DPI, 5mg.L^−1^), which is in agreement with our observation of a loss of function frameshift mutation in the alt-SDHC gene in this strain (position 78 in Figure 7B, S6 Table). On fluopyram-supplemented plate (18DPI, 5mg.L^−1^), a longer incubation was required to distinguish a wide range of growth phenotypes for the *alt-SDHC* containing strains, these varied from strong growth to no growth at all, including strains carrying a functional *alt-SDHC* gene. Under these conditions, some background growth started to become visible for isolates devoid of the *alt-SDHC* gene. However, only *alt-SDHC* containing strains displayed strong to moderate growth in the assay. Finally, for pydiflumetofen, in addition to an extended incubation, the concentration of the molecule needed to be reduced by 50 fold (18DPI, 0.1mg.L^−1^) to observe moderate growth with resistant controls 06STD024 and 07GB009 and with same isolates that displayed a strong growth on fluopyram-supplemented plates. Also for this compound, background growth started to become visible for a range of isolates not carrying *alt-SDHC.* This effect was even more pronounced than with fluopyram, suggesting that other genetic factors besides the presence of *alt-SDHC* are also relevant for baseline sensitivity differences to this molecule in the population.

**Figure 7.**
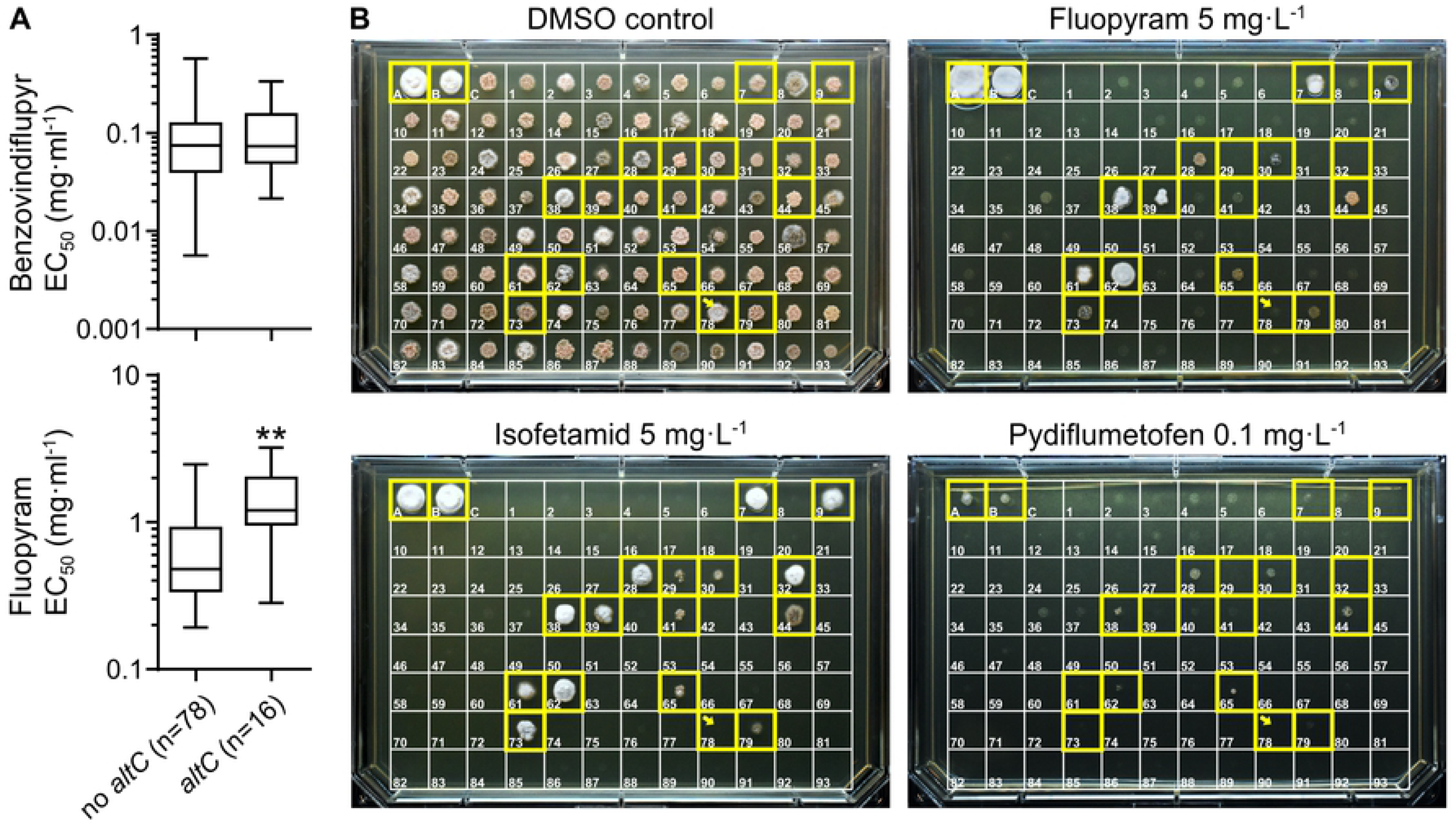
Resistance to SHA-SDHIs in *Z. tritici* EU populations. A. Box and whisker plot presenting EC_50_ sensitivity data of 93 *Z. tritici* isolates sampled in the EU in 2009. Sensitivity data are sorted by genotype according to the presence of the *alt*-*SDHC* gene. ** p value of 0.0029 in Welch’s corrected unpaired t-test. B. Solid agar growth of a collection of 93 *Z. tritici* isolates sampled in 2009 (same set as above). Individual strains from this collection are boxed and numbered 1 to 93. Boxes A and B correspond to reference resistant strains 06STD024 and 07GB009 respectively. Box C corresponds to IPO323 reference sensitive isolate. The yellow framed boxes correspond to strains carrying the *alt*-*SDHC* gene. Yellow arrow designates strain 09STIR20.1 (number 78) carrying a non-functional *alt-SDHC* (frameshift, S6 Table). Each individual strain was spotted onto AE agar plates (approx. 700 cells per spot) supplemented or not with isofetamid 5 mg.L^−1^, fluopyram 5 mg.L-1 and pydiflumetofen 0.1 mg.L^−1^. Plates were left to grow at 20°C in the dark and imaged at 10 DPI (DMSO control and isofetamid) or 18 DPI (fluopyram and pydiflumetofen).

These growth assay results are in good agreement with the different potency and resistance factors observed in SQR assays for SHA-SDHIs (Table 1). Isofetamid combined a high potency on WT-SQR and extremely high resistance factor on alt-SQR enzyme suggesting that even low quantities of the alt-SDHC protein could lead to a visible phenotype at high concentrations of the molecule. Fluopyram combined moderate potency on WT-SQR and high resistance factor on alt-SQR suggesting that at high concentration a good correlation would be maintained. Pydiflumetofen combined high potency on WT-SQR with low resistance factor on alt-SQR which suggested that at low concentrations a moderate to low correlation would be found.

A subset of eight isolates carrying the *alt-SDHC* gene was analyzed for their sensitivity to SDHIs, alt-SDHC expression and splicing patterns. Liquid culture growth sensitivity tests were performed to determine EC_50s_ for this set of isolates towards a wide range of commercial and research SDHIs (S5 Table). The results obtained for fluopyram are presented in figure 8B. The liquid culture sensitivity results are in good agreement with growth phenotype on solid agar at fixed concentration (Fig 8A, 8B). These experiments support a wide range of fluopyram SHA-SDHI resistance levels from 3 fold for 09STIR20.3 to 50 fold for 06STD024 and 07STGB009 (Figure 8B, S5 Table). We hypothesized that these differences in resistance levels among *alt-SDHC*-carrying isolates are driven by differences in its expression. Indeed, semi quantitative RT-PCR and hydrolysis probe RT-qPCRs revealed varying proportions of spliced and unspliced *alt-SDHC* mRNA across the range of tested isolates (Figure 8C and 8D). Total *alt-SDHC* mRNA correlated with increased splicing efficiency (Figure 8E). This efficiency ranged between not measurable for the most sensitive isolate 09STIR20.3, to 73% and 87 % for the most SHA-SDHI resistant isolates 07STGB009 and 06STD024 respectively. Moderately resistant isolates 09STF011 and 09STF112 displayed less of the spliced form, which represented 60% and 59% of total alt-SDHC mRNA respectively. Overall, the quantity of spliced *alt-SDHC* mRNA correlated with fluopyram resistance levels (Figure 8F). Interestingly, *SDHC* expression levels were concomitantly found to be the lowest in the most highly *alt-SDHC* expressing strains 06STD024 and 07STGB009 (Figure 8D), suggesting a possible link between the two.

**Figure 8.**
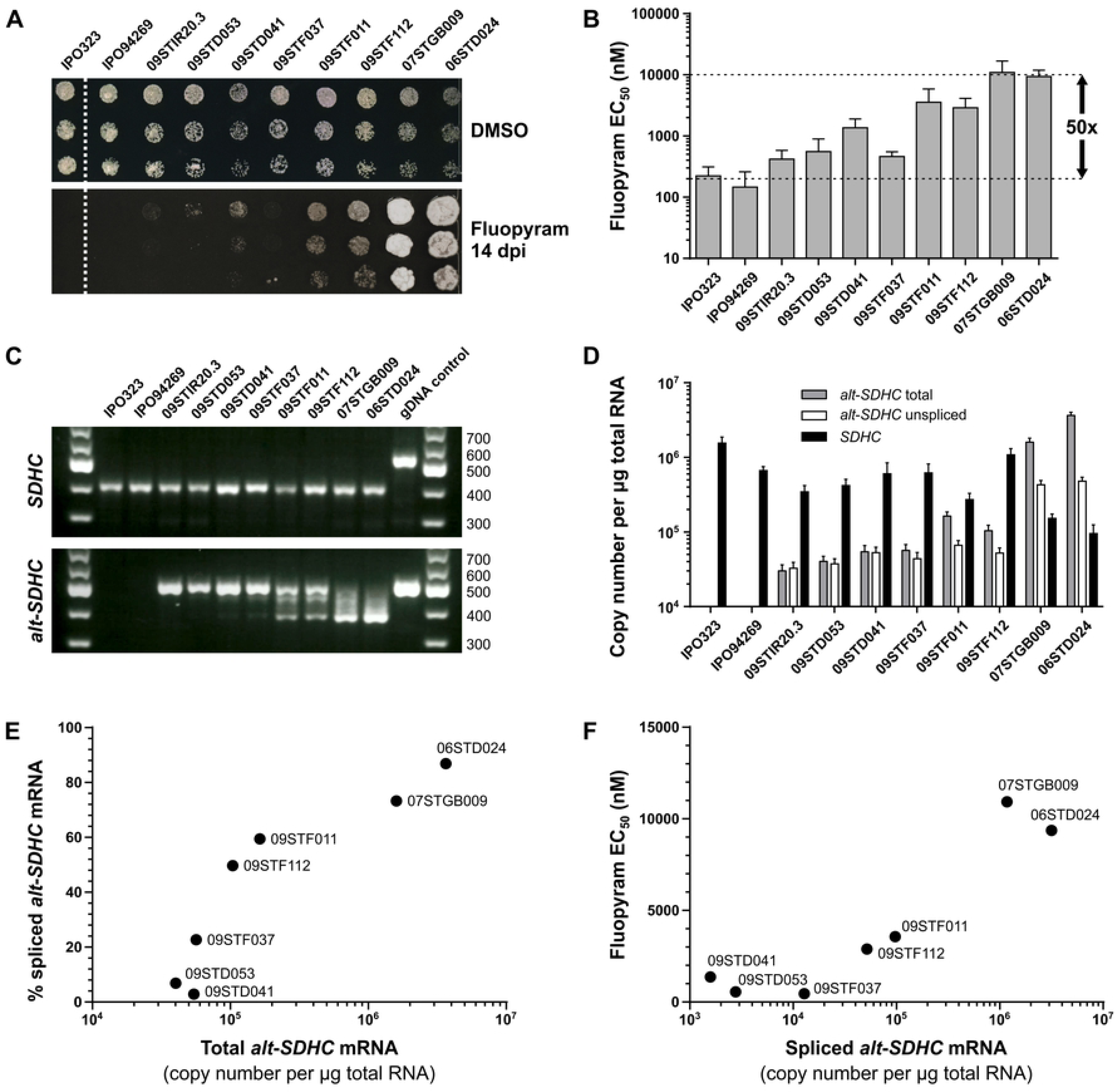
Fungicide sensitivity, gene expression and mRNA splicing in *alt*-*SDHC*-containing field isolates. A. Growth phenotypes of a collection of ten *Z. tritici* field isolates on solid AE agar supplemented or not with different SDHIs. Two control strains (IPO323 and IPO94269) lack the *alt*-*SDHC* gene whereas the other eight isolates (09STIR20.3, 09STD053, 09STD041, 09STF037, 09STF011, 09STF112, 07STGB009 and 06STD024) all carry the gene. B. Fluopyram liquid culture sensitivity results (similar set of strains as shown in A). C. Gel electrophoresis of RT-PCR products of *SDHC* and *alt*-*SDHC* (5’ regions encompassing 2 introns each). gDNA of strain 06STD024 was used as control. D. Absolute quantification by hydrolysis probe RT-qPCR of total *SDHC* mRNA, and of total and unspliced *alt*-*SDHC* mRNAs. E. Plot of total *alt*-*SDHC* mRNA for each isolate versus calculated percentage of spliced *alt*-*SDHC* mRNA. Results for strain 09STIR20.3 not shown (calculation leading to negative value). F. Plot of spliced *alt*-*SDHC* mRNA against fluopyram liquid culture sensitivity. Results for strain 09STIR20.3 not shown.

At the protein level, the total amount of mitochondrial SDHC proteins (alternative and core) ranged between 21 fmol in IPO94269 to 130 fmol in 07GB009 suggesting that the total amount of mitochondrial SQR protein varied across isolates (Table 3). Surprisingly, the alternative SDHC protein could be detected in all isolates carrying the gene, including the fully sensitive isolate 09STIR20.3 in which the alternative SDHC mRNA is very poorly expressed and for which splicing was not detected (Table 3, Figure 8C). Isolates 06STGB009 and 07STD024 which showed the strongest fluopyram resistance also displayed the highest amount of alternative SDHC (up to 120 fmol in 06STGB009). These high levels of alternative SDHC protein were associated with very low (0.4 fmol in 07STGB009) or undetectable (06STD024) amounts of the “core” SDHC protein. Moderately shifted isolates 09STF011 and 09STF112, in which balanced splicing of alt-SDHC mRNA was detected, also displayed a balanced abundance of both SDHC proteins from 38.6 to 57.7 fmol for alternative SDHC while the core SDHC protein was depleted but still detectable at 7.8 and 4.5 fmol respectively. Despite the differences in splicing efficiency, isolates 09STD041 and 09STF037 displayed very similar SDHC proteins quantities and ratios compared with 09STF011 and 09STF112. This was unexpected given the differences in RT-PCR suggesting lower quantities of the alt-SDHC protein should have been observed. Finally isolates 09STIR20.3 and 09STD053 for which no or very low splicing could be detected displayed amounts of core SDHC similar to WT isolates IPO323 or IPO94269. The alt-SDHC protein was detected at similar levels to other moderately or poorly shifted isolates in 09STD053 (48.6 fmol) and in much lower amounts in 09STIR20.3 for which no splicing of the alt-SDHC mRNA was detected (2 fmol).

**Table 3.** Quantification of the SDHC and altSDHC proteins in mitochondrial extracts of a panel of 10 field isolates by LC-MS/MS.

| Isolate | SDHC (fmol) | altSDHC (fmol) |
| --- | --- | --- |
| IPO323 | 41.9 ± 20.5 | nd* |
| IPO94269 | 21.1 ± 3.3 | nd* |
| 09STIR20.3 | 39.6 ± 20.9 | 2.0 ± 1.3 |
| 09STD053 | 27.1 ± 13.0 | 48.6 ± 17.0 |
| 09STD041 | 4.9 ± 4.9 | 46.3 ± 22.3 |
| 09STF037 | 5.1 ± 2.8 | 55.4 ± 3.2 |
| 09STF011 | 7.8 ± 3.5 | 38.6 ± 5.0 |
| 09STF112 | 4.6 ± 0.4 | 57.7 ± 9.7 |
| 07STGB009 | 0.4 ± 0.7 | 133.2 ± 15.9 |
| 06STD024 | nd* | 97.2 ± 47.0 |

Values presented correspond to the mean of 6 individual determinations on the same set of samples ± SD. \*not detected.

These data demonstrate the importance of expression levels and splicing efficiency of *alt-SDHC* mRNA in conferring the resistance phenotype. They also suggest that the alt-SDHC protein is more stable compared to the core SDHC in *Z. tritici* mitochondria, since very low expression of the functional spliced mRNA is sufficient for detection of the protein. Depletion of the core SDHC subunit, which is likely due to its replacement by alt-SDHC within the SQR enzyme, seems to correlate to the resistance phenotype.

### 2.7. Up-regulation of *alt-SDHC* gene expression in field isolates is associated with transposons insertions in the promoter region

The *alt-SDHC* locus of isolate 06STD024 differed from the corresponding locus in the 3D7 genome by the insertion of a large class II transposon (no cat element 8, 7kb) located 182 bp upstream of the *alt-SDHC* start codon (Figure 2D). Fragments encompassing the *alt-SDHC* gene as well as ∼1.5kb downstream and upstream sequences were amplified and sequenced in the eight isolates already characterized for fluopyram/SHA-SDHI resistance (Figure 8).

All isolates, except 07STGB009, displayed a similar *alt-SDHC* locus organization to 3D7 (Figure 9A). SNPs, insertions and deletions were detected in the intergenic region located between *alt-SDHC* and its 5’ neighboring gene (EMBL: ZT3D7_G4529, Figure 9A). The highest variation in this region corresponded to a 23bp deletion 80bp upstream of *alt-SDHC* start codon in moderately resistant isolate 09STF112.

**Figure 9.**
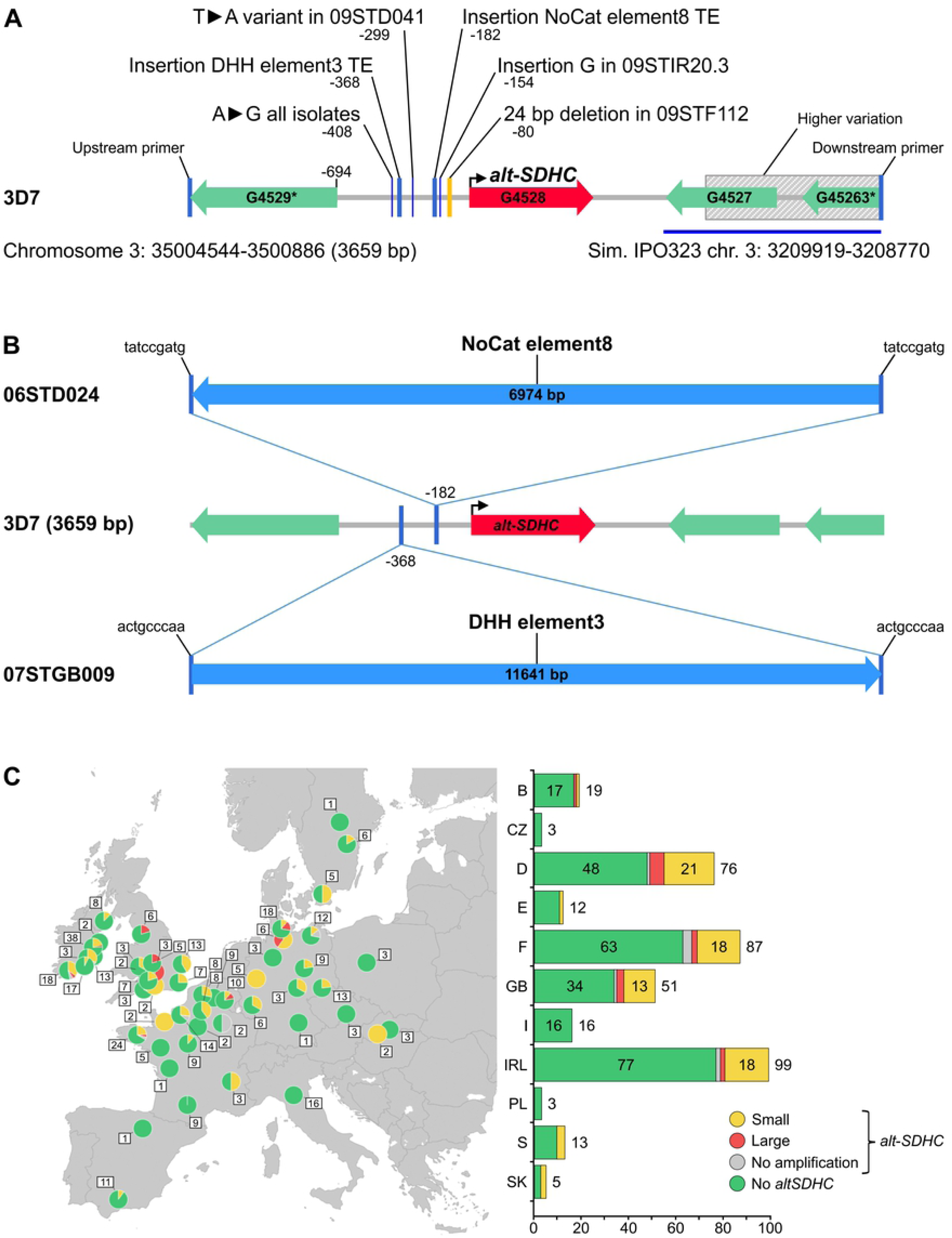
Structural variation at the *alt*-*SDHC* locus in European *Z. tritici* field isolates and populations. A. Structural overview of *alt*-*SDHC* locus variations in a set of sequenced isolates. Mutations are lined up to 3D7 genomic structure, only mutations located within the region between ZT3D7_G4529 start codon to the stop codon of the *alt*-*SDHC* gene are shown. Positions are numbered according to the *alt*-*SDHC* start codon (+1). Sequences have been deposited at NCBI under references MK067275-MK067282. B. Insertion of transposable elements in the promoter of *alt*-*SDHC* of highly resistant 06STD024 and 07STGB009 field isolates. Target site duplications of 9 bp are flanking each transposon insertion. C. European map with pie charts representing the 4 genotypes detected in *Zymoseptoria tritici* isolates collected in 2016. Green: *alt*-*SDHC* gene absent, grey: *alt*-*SDHC* gene present and no promoter amplification product, yellow: *alt*-*SDHC* gene present and promoter of classical size, red: *alt*-*SDHC* gene present and promoter of larger size. The total count of isolates for each sampling location is presented in white boxes. Right panel: Bar chart showing the total count of isolates of each type (similar color code) listed by countries.

Long range PCR was used to amplify a much larger fragment (12 kb) in the highly resistant isolate 07STGB009. In this isolate, we detected the insertion of a large class II DNA transposon of 11.6 kb in length and annotated DHH element 3 in the promoter region of the *alt-SDHC* gene (Figure 9B). This DNA transposon was also found at different genomic loci and in variable copy numbers among fully sequenced *Z. tritici* isolates. The transposon insertion site was located 368 bp upstream of the *alt-SDHC* start codon and at 195 bp upstream of the other transposon insertion site in the 06STD024 strain. In both cases, a 9 bp sequence of the *alt-SDHC* promoter was duplicated at the border of each transposon, suggesting the recent insertion of these transposons at these loci (target site duplication, Figure 9B). Overall the insertion of transposons in the promoter of *alt-SDHC* was only observed in the two highly resistant isolates 07STGB009 and 06STD024. This result suggested that the insertion of transposons in the promoter of *alt-SDHC* supports higher expression and better functional splicing of the gene.

### 2.8. Frequency of structural variants in *alt-SDHC* promoter in European *Z. tritici* populations

In order to explore the frequency of structural changes in the *alt-SDHC* promoter region at a population scale, a set of 145 strains carrying the *alt-SDHC* gene and collected during the years 2009, 2010 and 2016 in Europe was assessed using locus-specific primers (S5 Table). Within this set of 145 field isolates, amplification products of 2.4kb, similar to the expected size of 3D7 were obtained for 117 isolates (80.7%). Amplification products of larger sizes than 3D7, ranging between 3 and 20 kb, were found in 17 isolates (11.7%) and no amplification product was obtained for 11 isolates (7.5%). Insertion points were determined for 14 isolates displaying larger promoters (Table S8). The insertion points ranged between 173 and 1073bp upstream of the *alt-SDHC* start codon, suggesting a wide range of structural variations. A graphical overview of the results obtained for the 2016 population (n: 387 isolates) characterized for presence/absence of *alt-SDHC* gene and its promoter structure is presented on a graphical map of Europe (Figure 9C). The *alt-SDHC* gene is widely distributed across Europe, but was more frequently found in isolates from the United Kingdom, Ireland and Northern regions of Germany and France. Structural promoter variants corresponding to potential insertions of transposons were detected in isolates from Germany, United Kingdom, Ireland, France and Belgium. Although tested isolates with insertions upstream of the *alt-SDHC* gene being on average more resistant to fluopyram than isolates with no insertions (Figure S4), the difference between the two groups was not statistically significant.

## 3. Discussion

We demonstrated that a pre-existing SDHC paralog characterized by (i) its presence / absence and (ii) functional expression polymorphisms is responsible for standing sensitivity variation towards a particular class of SDHIs in the European *Z. tritici* population. Phylogenetic analysis showed that the *alt-SDHC* gene (*ZtSDHC3*) originates from an ancient duplication of an ancestor of *SDHC (ZtSDHC1)*. Another paralog, *ZtSDHC2*, is also present in all sequenced isolates. We initially did not consider this paralog as a potential SQR subunit because (i) the gene model was partly incorrect and lacked a mitochondrial targeting peptide and (ii) because the gene was not substantially expressed in either the IPO323 or 3D7 strains and in any of the conditions tested (S3 Figure), (36, 39). Functional explorations by reverse genetics will be required to assess whether this gene can perform a true SQR function. If this is the case, we propose that similarly to *alt-SDHC*, *ZtSDHC2* expression variants may exist in the population which will potentially further leverage our understanding of target-based SDHI sensitivity patterns in this pathogen.

*Rhynchosporium commune* was to our knowledge the only well described example of a plant pathogen carrying a dispensable target gene paralog responsible for standing fungicide sensitivity variation in populations. The presence of multiple paralogs of *CYP51*, the target of azole fungicides is common in ascomycetes. *R. commune* isolates can carry up to two functional *CYP51* paralogs, *CYP51A* and *CYP51B*. The *CYP51A* paralog is dispensable and was mostly absent in *R. commune* populations before azole adoption but re-emerged following the introduction of azole fungicides (40). The *R. commune CYP51A* paralog benefits from an azole-inducible regulation and confers a ten-fold sensitivity shift towards azole fungicides. A recent analysis of a global set of 400 *R. commune* isolates validated selection of the *CYP51A* gene at a global scale, and also the recent emergence of novel *CYP51A* variants carrying nonsynonymous substitutions, likely resulting from azole fungicides selection (41). The dispensable *CYP51A* paralog is therefore the main factor driving the sensitivity shift towards azole fungicides in *R. commune*.

Similarly to *R. commune CYP51* paralogs, the biological reasons for the emergence of multiple *SDHC* paralogs is unclear. It seems that duplication events of the *SDHC* gene have occurred multiple times throughout evolution in fungi, but the conservation (or loss) of the paralog(s) seems species-specific (S4 Table). By analogy to our findings one could suggest that the presence of paralogs could support standing resistance towards natural SDHIs. To our knowledge, few natural SDHIs have been reported so far. Siccanin (a metabolite from *Helminthosposium siccans*) and atpenins (metabolites from *Penicillium sp.*) can inhibit bacterial, fungal and mammalian SQRs through the Qp site (42–44). Metchnikowin an antifungal peptide of *Drosophila melanogaster*, was also recently found to bind to the *Fusarium graminearum* iron-sulfur subunit SDHB and to inhibit fungal SQR *in vitro* (45). Therefore, SDH inhibition by natural antifungal compounds as a driving force for the selection of *SDHC* paralogs in fungi cannot be totally ruled out. The *Z. tritici SDHC* paralog situation could therefore result from a specific acquired resistance profile towards natural products synthesized by competing species, similar to bacterial antibiotics resistance genes which have been shown to be of ancient origin because they evolved to resist pre-existing natural products (46).

The core role of the succinate dehydrogenase step of the TCA cycle in driving primary metabolism and consequently growth and secondary metabolism may rather suggest that *SDHC* paralogs would permit a controlled production of hybrid SQR enzymes. This could have subtle effects on growth phases during pathogenicity and may carry an advantage leading to their maintenance within populations, particularly if the gene is under developmental regulation and leads to a SQR enzyme of differing efficiency as found in parasitic nematodes (47, 48) or plants (49). In yeast, one paralog of the SDHC subunit and one paralog of the SDHD subunit were shown to lead to hybrid functional SQR enzymes which, although less active, may play an adaptive role in restrictive environmental conditions (50). Similarly to mutants of the classical SDHC and SDHD subunits, the hybrid SQRs conferred very distinct “metabotypes” which also correlated with different growth yield (50, 51). Interestingly, the yeast *SDHC* paralog may also carry additional function(s) as the protein was found in a subcomplex with Tim18p as part of the TIM22 inner membrane translocase (52). Altogether this suggests that *SDHC* and its paralogs may possibly functionally overlap for a range of functions.

Based on its complex regulation and partial splicing it will be interesting to determine whether *alt-SDHC* displays particular expression or splicing patterns during *in planta* infection. Within our panel of eight isolates specifically chosen for covering a range of resistance phenotypes, expression levels of spliced *alt-SDHC* and in particular the amount of spliced *alt-SDHC* positively correlated with the SHA-sensitivity shift (Figure 8). The main factor limiting SHA resistance was the incomplete splicing of the mRNA which was clear in all isolates and appeared the least effective in low expressing non-shifted isolates. The replacement of the SDHC subunit within the functional SQR complex was associated with disappearance of the SDHC protein from the mitochondria suggesting degradation of the non-SQR-integrated polypeptide. This effect was particularly clear within the highly shifted, highly expressing strains in which the core SDHC protein was significantly depleted. Conversely, the alt-SDHC protein seemed less prone to degradation since low levels of the protein were still observed in strains expressing low levels of the gene, including strain 09STIR20.3 for which spliced *alt-SDHC* mRNA was not detected by RT-PCR (Table 3, Figure 8). If not an artefact, this differential degradation pattern suggests that either the alt-SDHC polypeptide outcompetes SDHC protein for inclusion within the functional SQR or that the protein gets integrated within another complex of the IMM of another function. Alternatively, a less effective scavenging of the complex-free alt-SDHC polypeptide could lead to the same effect (53).

Interestingly, limited sequence variation was observed within the *alt-SDHC* gene, which highly contrasts with very high sequence variability in the *Z. tritici ZtSDHC1* gene. This suggests that the two genes undergo very different selection pressures. We hypothesize that the higher evolutionary pressure on *ZtSDHC1* is linked to its principal SQR function. The membranous SDHC and SDHD subunits are the least evolutionarily conserved SQR subunits (4) and in *Z. tritici* populations both genes show high sequence variation. To form the SQR enzyme, the SDHC and SDHD subunits associate as an integral membrane heterodimer of the IMM and serve as membrane anchors to the mitochondrial matrix SDHA/B catalytic dimer. It is conceivable that *SDHC* and *SDHD* variant combinations may not all be similarly favorable to the translation and scaffolding of functional SQR. Natural variations within the core *SDHC* (*ZtSDHC1*) could have a biological impact in regulating SQR amounts and efficiency, in particular non-synonymous variations such as the highly frequent C_N33T, N34T allele within the transit peptide may possibly impact mitochondrial import efficiency whereas the multiple synonymous mutations found in the gene may lead to differential translation efficiencies. Variations of the SDHC and SDHD genes could therefore represent a means to differentially regulate SQR-associated developmental or metabolic traits that are important for pathogenicity. So it can be envisaged that either one or the two paralogs of *SDHC*, the conserved *ZtSDHC2* and the dispensable *alt-SDHC* have a biological significance in building alternative hybrid SQR enzymes and regulating developmental growth and metabolism at particular stages of the infection. Interestingly, *in planta* assays with isogenic lines carrying SDHI resistance-conferring mutations showed increased necrosis on a wheat cultivar suggesting a link between functional efficiency of the SQR enzyme and virulence (14).

An in depth exploration of haplotype networks of the *ZtSDH* genes is under investigation and may provide further support to this hypothesis. In general, the higher conservation of *alt-SDHC* combined with its higher stability within mitochondrial membranes could suggest an opportunistic integration within the SQR complex. The *alt-SDHC* may represent an independently evolved *SDHC* paralog which could have undergone convergent functional evolution after initial divergence.

Fluopyram is currently the only SHA-SDHI molecule registered for STB control in Europe. Given the complete lack of control observed *in planta* for resistant isolates we expect that poor efficacy and strong selection would result from applications of the solo compound (21). However, fluopyram is sold in a mixture with another SDHI (bixafen), and an azole (prothioconazole) which are both not affected by the mechanism. Novel highly active SHA-type SDHI molecules such as pydiflumetofen, which are intrinsically more active on *Z. tritici* compared to current SDHIs, will likely also exert a selection for strains carrying the alternative *SDHC* paralog. However, unlike *CYP51*, the two *SDHC* paralogs (i) display differential sensitivity towards compounds of the same mode of action and (ii) compete for inclusion into the functional SQR which partially limits the sensitivity shift. Therefore, we may expect that such mechanisms would be outcompeted by classical target mutations of the core genes such as the C-H152R mutation, conferring superior sensitivity shifts and cross-resistance towards all classes of SDHIs (17, 18). Using a GM approach we artificially generated isolates only carrying the alt-SQR enzyme. This enabled us to compare SHA-SDHIs of different structures on the two types of SQR enzymes and to gather rational understanding of the structural features maximizing potency and minimizing cross-resistance. The pure alt-SQR GM isolates represent a worst case scenario for this paralog-mediated resistance mechanism. We observed that despite a superior sensitivity shift of the GM isolate (KO-*SDHC*) compared to the original isolate *in vivo*, the dose of pydiflumetofen required for full *in planta* control was similar to that of other SDHIs on the market for controlling wild type isolates. This result has high practical relevance since it supports the use of a robust rate for the novel SHA-type SDHIs for an effective control of the whole population including strains carrying the *alt-SDHC* gene. As such our findings explain baseline variation for SHA-SDHIs and will enable tracking for potential evidence of selection such as an increased occurrence of strains carrying the gene or the potential emergence of mutated forms of the alternative protein.

The decreasing cost of population-wide genome sequencing and the widespread adoption of genome-wide association studies (GWAS), should facilitate the identification of the molecular factors involved in baseline fungicide sensitivity (33, 54). Since the increased natural variation encompassed by paralogs provide a source of direct adaptation to natural compounds or xenobiotics it is likely that population genomics will enable the discovery of many more instances of dispensable paralogs of fungicide targets or detoxification genes. For agrochemical research, population variation is a major challenge that needs to be addressed and an in depth understanding of the molecular factors involved represents a real opportunity for more accurate chemical design of future solutions.

## 4. Materials and Methods

### 4.1. Strains, media and culture conditions

All *Z. tritici* strains were isolated from infected wheat leaves collected during Syngenta European monitoring following already described procedures (22). The reference strains IPO323 and IPO94269 were kindly provided by Gert H.J. Kema (Wageningen University, NL). The isolates were inoculated from stocks stored in liquid nitrogen onto solid V8 agar at 18°C for 5 days (55). Fresh cells were harvested from these plates and used as an inoculum for all experiments. The following media were used throughout: TSM40 (4 g.L^−1^ glucose, 10 g.L^−1^ malt extract, 4 g.L^−1^ yeast extract, pH 7.0); AE medium (56); induction medium (IM) (55), YPD (10 g.L^−1^ yeast extract, 20 g.L^−1^ peptone, 20 g.L^−1^ glucose). DH5α, TOP10 or DB3.1 cells (Invitrogen) were used for the maintenance of plasmids in *Escherichia coli*. *Agrobacterium tumefaciens* strain EHA105 (57, 58), was used for *A. tumefaciens* mediated transformation (ATMT) following procedures described in (55).

### 4.2. Liquid culture assays for fungicide sensitivity determination

Pre-culture of the inoculum and fungicide sensitivity tests were performed following previously described procedures (14). Different ranges of inhibitor concentrations were used for population monitoring and for the detailed phenotyping of a selected set of individual field isolates and genetically modified strains. For fungicide sensitivity monitoring of European field populations, final inhibitors concentrations were between 100 mg.L^−1^ and 0.0001 mg.L^−1^ with uniform 10x dilution steps (7 inhibitor concentrations + DMSO control). For refined sensitivity analysis of a smaller panel of isolates final inhibitor concentrations ranged between 0.5 mM and 0.47 nM with uniform 4x dilution factor steps (11 inhibitor concentrations+ DMSO control).

### 4.3. Mapping population generation and resistance mapping

Mating type determinations were performed using PCR markers described in (59). 06D024 × IPO323 and 07GB009 × IPO94269 crosses were performed as previously described in (60). Single ascospore progeny isolates were collected and groups of 234 and 96 isolates were obtained respectively. Fluopyram resistance inheritance was determined by spotting 2µl of 2.10^6^ cells.ml^−1^ onto AE agar supplemented or not with fluopyram 10 mg.L^−1^. DNA extraction were performed using fresh culture grown on V8-agar plates (5 days, 18°C in the dark), approximatively 100 mg of fresh cells were collected with an inoculation loop and processed to DNA extraction using the DNeasy 96 Plant Kit (Qiagen) and following provider’s instructions. For pool sequencing, 2 µg of DNA for each pool was sheared to an average fragment size of 340 base pairs using a Covaris S220 focused-ultrasonicator (Covaris, Inc., Woburn, Massachusetts, USA). The samples were then cleaned using DNAClean XP (Beckman Coulter Life Sciences, Indianapolis, Indiana, USA). Sequencing libraries were prepared from the sheared DNA using the NEBNext® DNA library prep kit for Illumina (New England BioLabs, Ipswich, Massachusetts, USA). Size selection was performed using an E-gel precast agarose system. Each sample was run in three lanes of an Illumina Genome Analyzer II (Illumina, San Diego, California, USA) in a 36 cycle paired end run. Total sequence yield was 2.7 gigabases for the resistant pool sample and 3.3 gigabases for the susceptible pool. Sequence reads were aligned to the JGI *M. graminicola* v2.0 assembly, using gsnap (61) and uniquely aligning reads were used to call variants with the Alpheus pipeline (62), with filtering criteria requiring at least 2 reads having average base quality of ≥ 20 with an allele frequency within the sample of ≥ 0.2. Differences in allele frequencies between the pools were then used to determine the putative genomic location of the causative variants. All PCR-based genotyping assays (SSR, CAPS) were run on individual genomic DNA using GoTaq DNA polymerase, at recommended temperature and cycling parameters and using oligonucleotides and enzymes listed in S7 and S1 Tables respectively.

### 4.4. Phylogenic analysis of fungal SDHC proteins

Orthologs for SHDC1 were retrieved using ENSEMBL ortholog/paralog prediction where available (63). For *ZtSDHC2* and *ZtSDHC3* or genomes not ortholog mapped in ENSEMBL a reciprocal BLAST was performed to identify homologous sequences. All retrieved sequences were run through TargetP analysis (34) and only mitochondrially targeted sequences were retained. Sequences were aligned using Clustal-omega with default settings (64). A tree was drawn using PhyML for amino acid sequences using the best of NNI and SPR as the tree topology search operation (LogLk = −21960.63386) (65). The tree was visualized using iTOL (66).

### 4.5. PCR methods and Sanger sequencing

All oligonucleotides were purchased from Microsynth AG (Balgach, 175 Switzerland). PCR primers used to amplify sequences for CAPS/SSR markers, Sanger sequencing or clonings are listed in S3 Dataset. PCR products for cloning or direct Sanger sequencing were obtained using the Phusion Hot Start II High-Fidelity DNA Polymerase (ThermoFisher Scientific, F549L). For the long PCR products required to characterize promoter inserts, LongAmp Taq DNA Polymerase (NEB, M0323S) was used. PCR products for classical genotypings such as CAPS markers or SSR analysis were amplified using GoTaq® G2 Hot Start Polymerase (Promega, M7405). Each PCR was performed according to the conditions recommended by the respective manufacturers. Sanger sequencing was done at Microsynth AG (Applied Biosystems 3130 Genetic Analyzer). Pyrosequencing was performed on a PyroMark Q96 ID (Biotage/QIAGEN).

DNA was extracted using the DNeasy 96 Plant Kit (Qiagen) following provider’s instructions. For the sequencing of large promoter insertions, the large fragments were cloned into TOPO vectors and a primer walking procedure applied at Microsynth AG (Balgach, 175 Switzerland).

### 4.6. Growth tests on solid agar at discriminatory fungicide concentrations

A large scale spotting assay of 96 isolates (Figure 7) was performed using the V&P 96 floating pin tool VP408FP6 (V&P Scientific), equipped with flat tip FP6 pins of 1.58 mm diameter (resulting in approx. 0.4 µl transfer volume). Source cultures for cell spotting were grown in 100 µl YPD liquid medium in a 96 well flat bottom plate (Corning, 3370) at 18 °C for 11 days (average cell density of 3.5·10^6^/ml), then diluted 1:2x in fresh YPD medium and incubated for another 1.5 h before transfer with the pin tool (approx. 700 cells per spot) onto AE agar with or without fungicide. Plates were incubated at 20 °C in the dark for up to 18 days. Smaller scale spotting assays (Figure 4 and Figure 8) were performed using 5 days V8-agar plates inoculums adjusted to 2.10^6^ spores.mL^−1^ in water and diluted in steps of 3. 2µL of spore suspension was spotted on the plates, and the plates were incubated shielded from light at 21°C for 6 days.

### 4.7. *In planta* fungicide dose response

Wheat (*Triticum aestivum*) variety Riband was grown in pots (d = 6.5cm), at a density of 4 plants per pot and treated with the growth regulator CCC (Chlorcholinchlorid; Chlormequat; 5 ml / pot, 0.4% solution) 4 days after sowing. Wheat plantlets were maintained in a climatic room at 18°C, 60% humidity and under a 12h light regime (high intensity). Fungicide applications were performed on 14 days old plantlets for which leaf 2 is the fully expanded target leaf. Fungicide treatments were performed using a custom-made track sprayer adjusted at 200L.ha^−1^ (Nozzle: Lechler, orange LU90-01). The fungicides used were Solatenol^TM^ EC100 (Elatus Plus, benzovindyflupyr), Isopyrazam EC125 (Seguris Flexi or Reflect), Adepidyn^TM^ EC100 (research formulation of pydiflumetofen). *Z. tritici* infections were performed using a Devilbis airbrush (spray of about 150ml.m^−2^) one day after fungicide application and using an inoculum grown on V8-agar adjusted to 1.8.10^6^ spores.ml^−1^ in 0.05% Tween20 in MQ water. Inoculated plants were initially incubated for 72h under reduced light conditions and high humidity using towel-covered Plexiglas hoods in a climatic chamber set to 21°C/19°C day/night alternations, 80% humidity and a 14h light regime. The Plexiglas hoods were then removed until evaluation. Plants were fertilized once per week and disease evaluation performed based on disease coverage on the second leaf approximately 16-19 days after infection, once untreated plants reached 75-90% disease coverage. Each fungicide was tested at several rates to produce dose responses. There were 3 pots (4 plants each) per fungicide rate and isolate, the whole experiment was repeated 4 times. *In planta* EC_50_ were calculated using the software GraphPad Prism v6.08.

### 4.8. Production of *Z. tritici* transformants

The multisite binary pNOV2114_gateway and pNOV2114 Hyg _gateway (3-way) vectors were used to generate the different transformation constructs (14). To generate the *SDHC* and *alt-SDHC* KO mutants, 5’ upstream regions of 1000bp and 2074 bp and 3’ downstream regions of 914bp and 1313bp for *SDHC (ZtSDHC1)* and *alt-SDHC (ZtSDHC3)* respectively were PCR-amplified from genomic DNA of IPO323 or 06STD024 strains and the fragments cloned by BP cloning using Gateway™ BP Clonase II Enzyme Mix (Invitrogen) into pDONR-P4-P1R (upstream regions) or pDONR-P2R-P3 (downstream regions) (S3 dataset for oligos). These 5’ and 3’ gene locus-paired entry plasmids were then combined with the pENTR221-TrpChyg described previously (55) and pNOV2114_gateway for multisite gateway LR cloning using Gateway™ LR Clonase II Plus enzyme Mix (Invitrogen) following provider’s instructions. The final pNOV2114 KO-SDHC and pNOV2114 KO-alt-SDHC binary plasmids carry a hygromycin resistance cassette flanked by 5’ and 3’ upstream regions of the *SDHC* and *alt-SDHC* genes respectively.

For generating expression constructs under the control of a tetracyclin-repressible promoter, the plasmid pMF2-4h (67) was modified by removal of the hygromycin resistance cassette after digestion by *NotI* and recircularization of the plasmid to generate pMF2-4h^−^. The fragment containing the full Tet repressor expression cassette followed by operator sequences fused to *MfaI* minimal promoter was PCR amplified from re-circularized pMF2-4h^−^ plasmid and cloned by gateway cloning into pDONR_P4P1R using oligos listed in S3 dataset.

To generate the *alt-SDHC* expression plasmids, the *alt-SDHC* gene of 06STD024 was amplified from the genome and cloned into pDONRZeo by gateway cloning to generate pENTRZeo-alt-SDHC. A variant of this plasmid (pENTRZeo-alt-SDHC_I78A) encoding the I78A variant of *alt-SDHC* was obtained by site-directed mutagenesis using QuikChange® II Site Directed Mutagenesis kit (Stratagene) following provider’s instructions and oligos listed in S3 dataset. The Tetoff promoter region from plasmid pMF2-4h (67) was sub-cloned into pDONR221 using oligos listed in the S3 Dataset. These entry plasmids were combined with pENTR_TrpCterm and pNOV2114 Hyg_gateway plasmids (55) to generate the pNOV2114_Tetoff_alt-SDHC_TrpCterm and pNOV2114_Tetoff_alt-SDHC^I78A^_TrpCterm binary vectors used for transformation of IPO323.

All entry and subsequent binary plasmids were validated by Sanger sequencing of the cloned fragment before transfer to *A. tumefaciens. Z. tritici* transformation was performed as described previously (55). *Z.tritici* transformants were validated by PCR using primer combinations enabling the validation of successful gene deletion events for the KOs mutants or the completeness of the transformation cassette for the ectopic expression mutants.

### 4.9. Quantitative Real-Time PCR and semi-quantitative PCR

To produce the RNA samples, field isolates and transformants were initially inoculated on V8-agar plates and left to grow for 4 days, 25 ml TSM40 liquid cultures in 100 ml round bottom Erlenmeyer flasks were then initiated using 10 µl inoculation loops. The flasks were incubated at 20 °C, 160 rpm, for 4 days before cells were harvested by filtration using a tissue filter and ground in liquid nitrogen using mortar and pestle. For RNA extraction, 50 mg of the powdered material was processed with the RNeasy Plant Mini kit (Qiagen, 74904) according to the manual and including an on-column DNase I digestion (Qiagen, 79254). A second DNase digestion was performed on the eluates, followed by purification using the same

RNeasy Plant Mini kit. RNA yield and integrity was determined on an Agilent 2100 Bioanalyzer System and the absence of residual genomic DNA in the samples was verified by PCR, using primers specific for the β-tubulin (*TUB1*) gene (S3 Dataset) and the cycling protocol described below for semi-quantitative PCR. The High Capacity cDNA Reverse Transcription kit (Applied Biosystems, 4368814) was used for reverse transcription of 2 µg of total RNA per sample, using the RT Random Primers provided in the kit and according to the manufacturer’s instructions.

Semi-quantitative PCR was performed using GoTaq G2 Flexi DNA Polymerase (Promega, M7805) and the PCR primers listed in (S3 Dataset). *SDHC* and *alt-SDHC* from field isolates and reference strains were amplified from undiluted cDNA, whereas cDNA for detection of the β-tubulin sequence *TUB1* and of the *alt-SDHC* expression strain (samples pTet::*altC* and pTet::*altC* + Dox) were diluted 1:3x in water before use. Genomic DNA of isolate 06STD024 was included as control (carrying un-spliced template sequences for all three targets). The PCR program was: Initial denaturation for 2 min at 95 °C, followed by 40 cycles of denaturation at 95 °C for 30 s, primer annealing at 54 °C for 30 s and extension at 72 °C for 34, followed by a final incubation for 5 min at 72 °C.

Quantitative Real-Time PCRs were performed with all four targets in a multiplexed reaction using hydrolysis probes carrying different fluorophores and quenchers listed in S3 Dataset. The binding sites of qPCR oligonucleotides within *SDHC1* and *alt-SDHC* are shown in S5 Figure. Primers were used at 900 nM and hydrolysis probes at 200 nM final concentration in 20 µl multiplexed qPCRs with KAPA Probe Force qPCR Master Mix 2x (Kapa Biosystems, KK4301) and 5 µl template DNA per well. The cDNA preparations were diluted 1:9x in DEPC-treated water immediately before the experiment. RNA (No-Reverse-Transcription reaction controls) of the same samples were also tested in two separate runs using the corresponding plate layout.

To enable absolute quantification, a reference plasmid was generated by the sequential cloning of the coding sequences of *ZtSDHC1*, *TUB1* (both amplified from cDNA of IPO323), and *alt-SDHC* (from gDNA of 06STD024) using a GENEART Seamless Cloning and Assembly Kit (Invitrogen, A13288) and PCR oligos listed in S3 Dataset. The cloned fragments encompass the binding sites of the qPCR oligonucleotides. The resulting plasmid pUC19_cSDHC_gAlt-SDHC_cTUB1 (calculated molecular weight: 3200964.1 Da) was used to generate standard curves for both calculation of primer efficiencies and the absolute quantification of *ZtSDHC1* and *alt-SDHC* copy numbers. A serial 1:6x dilution of pUC19_cSDHC1_gAlt-SDHC_cTUB1 was made in 4 replicates, with a starting concentration of 4 pg/µl (resulting in 1 pg/µl or 20 pg total in the final reaction mix, 1 pg equals 188131 molecules based on the calculated molecular weight of 3200964.1 Da).

All 12 cDNA samples, a no-template RT reaction control and the reference plasmid dilution series were run on the same 96-well assay plate, with 4 technical replications per plate and the run was repeated on a duplicate plate. The qPCR was performed on a CFX96 Real-Time System on top of a C1000 Touch Thermal Cycler (Bio-Rad), and analyzed using the CFX Manager 3.0 software (Bio-Rad). The PCR program was: Initial denaturation for 3 min at 98 °C, followed by 45 cycles of denaturation at 95 °C for 10 s and combined annealing/extension for 20 s at 60 °C with subsequent plate reading. Assay results were exported to RDML format (S4 Dataset). For relative expression level comparisons of *ZtSDHC1* and *alt-SDHC*_total/un-spliced the Starting Quantity (SQ) values of individual wells were used to calculate the respective copy numbers using the reference plasmid standard curves. Statistical analysis was then performed on the 8 technical replicate values from individual wells.

### 4.10. Mitochondria isolation and enzyme assays

Biomass production, mitochondrial extraction and purification were performed as described in (14). Succinate: ubiquinone/DCPIP sensitivity tests were performed as described in (14) with minor modification. The different mitochondrial suspensions were adjusted to similar initial velocity (1 OD_595nm_ hour^−1^) and inhibitor concentrations ranged between 0.047nM and 50µM with uniform 4x dilution steps (11 concentrations + DMSO control). Calculated absorbance slopes (OD/min) were used for IC50 calculations using GraphPad Prism 6.07 software non-linear curve fitting against log inhibitor concentrations.

### 4.11. Sample preparation for SDHC and alt-SDHC protein quantitation

Protein from mitochondrial extracts was precipitated using trichloroacetic acid/acetone. After resuspension under denaturing conditions, the total protein concentration was estimated using a Bradford assay (68). An aliquot of 25 µg protein from each sample was separated on a 10% NuPAGE gel (Life Technologies). Gels were stained with colloidal Coomassie blue, and a gel region (10-25 kDa) from each lane was excised for trypsin digestion. In-gel digestion was carried out using a published protocol (69). After digestion, peptide samples were dried using a centrifugal evaporator, and re-suspended in LC-MS/MS sample buffer containing 3% acetonitrile, 0.1% formic acid, 100 femtomole (fmol) per microliter isotopically labelled internal peptide standards (JPT Peptide Technologies GmbH, Berlin, Germany). Peptide sequences and isotopic labelling information can be found in S7 Table. The peptides used for the LC-MS/MS analysis were chosen based on sequence uniqueness in the *Z. tritici* proteome. The peptides had also been identified in a separate proteomic analysis of mitochondrial extracts. Four technical replicates for each strain were prepared for LC-MS/MS analysis.

### 4.12. Multiple reaction monitoring LC-MS/MS analysis

LC-MS/MS analysis was done using a TSQ Vantage triple quadrupole mass spectrometer equipped with a nano-electrospray source (Thermo Fisher Scientific, Waltham, MA, USA) and coupled to an Ultimate/ Switchos split-flow LC system (Dionex, Thermo Scientific). A volume of 2.5 μl of each peptide sample was injected into the system. Peptides were separated on a Picotip column (75 µm, 15 cm column packed with 5 µm C18 particles; Nikkyo Technos Co., Ltd. Japan). Gradient elution was performed using 0.1% formic acid in water as solvent A and 99.9% acetonitrile/0.1% formic acid as solvent B. Gradient length was 30 min, from 3 to 40% solvent B. The flow rate was 300 nL per min. The TSQ Vantage instrument was operated with a capillary temperature of 275°C and spray voltage set to 1.7 kV. The data were acquired in positive scan mode with the collision gas set to 1.5 mTorr. The Q1 and Q3 peak widths (FWHM) were set to 0.2 u and 0.7 u, respectively. The cycle time was set to 5 seconds. No retention time scheduling for the two peptides was used. The list of monitored transitions and collision energy settings can be found in S8 Table. The run order of samples was randomised. The mass spectrometry raw files were imported into Skyline v1.2 software (University of Washington, USA). Integrated peak areas were exported to Microsoft Excel. The amount of SDHC and alt-SDHC protein in femtomole (fmol) was calculated based on the peak area for the endogenous peptide and the corresponding isotopically labelled internal peptide standard.

### 4.13. Homology model, docking simulations and conformational analysis

The homology model for *Z. tritici* WT-SQR with the “core” SDHC was generated as described in (14). The homology model of the alt-SQR carrying the alternative SDHC subunit was generated following a similar procedure. Isofetamid, pydiflumetofen and compounds 1-3, were manually docked into the *Z. tritici* SQR Qp binding site. Interactions of key residues were determined through pharmacophore elucidation (14). In a second step the protein ligand complexes were minimized using Moloc MAB force field (70), allowing full flexibility for the ligands while keeping the *Z. tritici* SQR protein rigid. For pydiflumetofen conformational analysis, a diverse set of 30 conformations were generated with the CCDC conformer generator (71). Each conformation generated by the CCDC conformer generator was optimized with the M06L DFT (72) functional method and 6-31G(d) as basis set within Gaussian09 (73). Additional parameters that were used: scrf=(iefpcm,solvent=water). Conformations have been evaluated based on the calculated DFT energy.

## 5. Acknowledgments

We would like to warmly thank Els Verstappen (Wageningen University) for her kind support with *Z. tritici* crossings, Dominique Edel (Syngenta) for her support with liquid culture cross resistance tests and Andrew Farmer (Syngenta) for his support with BSA analysis. Finally we would like to thank all the people who kindly reviewed this manuscript.

## 8. Supplementary material

**S1 Figure. Structure of carboxamides SDHIs molecules used in the study.**

Shaded grey area represents the SHA cross-resistance group. Schematic view of a typical SHA compound is shown in the bottom right corner.

**S2 Figure. Phylogenic tree of fungal SDHC proteins.**

Tree generated using PhyML and visualized using iTOL (see material and methods). ZtSDHC1-3 paralogs are highlighted in yellow.

**S3 Figure. Box Plot of IPO323 time course expression of SDH-encoding genes**

Expression data inferred from RNAseq (Rudd et al., 2015).

**S4 Figure. Box plot of fluopyram sensitivity of 133 *Z. tritici* isolates collected in Europe in 2016.**

Isolates were grouped by genotyping based on the detection of the *alt*-*SDHC* gene and *alt*-*SDHC*-promoter insertions. *ns: No significant difference in student t-test.

**S5 Figure. Schematic view of oligo positioning for *ZtSDHC1* and *ZtSDHC3* Taqman RT-qPCR assays.**

Exons are shown as blue arrows and introns as grey bars, labelled hydrolysis probes are shown in red, forward and reverse PCR oligos are shown as black arrows. Oligonucleotides sequences and probe details are shown in S3 Dataset.

**S1 Table. IPO323×06STD024 progeny genotyping results inferred from CAPS and SSR assays.**

**S2 Table. IPO323 genes within the final 16 kb mapping window.**

**S3 Table. 07STGB009×IPO94269 progeny genotyping results inferred from CAPS and SSR assays**

**S4 Table. Count of SDHC paralogs per species**

**S5 Table. Liquid culture SDHIs sensitivity for the panel of *Zymoseptoria tritici* field and genetically modified strains referred in the study.**

Mean: mean EC_50_ in nM, SEM: standard error of the means, N: number of individual determinations.

**S6 Table. Overview of *alt-SDHC* sequencing and promoter PCR results for a panel of 154 isolates carrying the gene and collected in Europe in 2009, 2010, 2011 and 2016.**

**S7 Table. Internal peptide standards used in the LC-MS/MS assay to quantify SDHC and alt-SDHC proteins.**

**S8 Table. Monitored transitions in LC-MS/MS assay for quantifying SDHC and alt-SDHC proteins.**

The assay used a multiple reaction monitoring approach on a TSQ Vantage triple quadrupole mass spectrometer.

**S1 Dataset. Core *SDHB*, *SDHC* and *SDHD* genes sequencing results for a set of *Z. tritici* field isolates referred in the study**

Base count according to first codon.

**S2 Dataset. Pool seq genotyping results and mapping interval**

**S3 Dataset. Oligonucleotides used in the study**

**S4 Dataset. q-RTPCR results files in RDML format (zip)**

